# Nemo-like kinase disrupts nuclear import and drives TDP43 mislocalization in ALS

**DOI:** 10.1101/2025.01.27.635090

**Authors:** Michael E. Bekier, Emile Pinarbasi, Jack J. Mesojedec, Layla Ghaffari, Martina de Majo, Erik Ullian, Mark Koontz, Sarah Coleman, Xingli Li, Elizabeth M. H. Tank, Jacob Waksmacki, Sami Barmada

**Author notes:** Correspondence to Sami Barmada: 109 Zina Pitcher Place, Ann Arbor, MI, 48019. contributed equally (Michael E. Bekier, Emile Pinarbasi).

## Abstract

Cytoplasmic TDP43 mislocalization and aggregation are pathological hallmarks of amyotrophic lateral sclerosis (ALS). However, the initial cellular insults that lead to TDP43 mislocalization remain unclear. In this study, we demonstrate that Nemo-like kinase (NLK)—a proline-directed serine/threonine kinase—promotes the mislocalization of TDP43 and other RNA-binding proteins by disrupting nuclear import. NLK levels are selectively elevated in neurons exhibiting TDP43 mislocalization in ALS patient tissues, while genetic reduction of *NLK* reduces toxicity in human neuron models of ALS. Our findings suggest that NLK is a promising therapeutic target for neurodegenerative diseases.

## Introduction

Amyotrophic lateral sclerosis (ALS) is a fatal and progressive neurodegenerative disease distinguished by loss of motor neurons in the cortex and spinal cord (1). The clinical presentation of ALS is heterogeneous, depending in large part upon the neuroanatomical pathways involved. Despite this, a single pathological hallmark — mislocalization and aggregation of the RNA binding protein TDP43 — is found in >95% of individuals (2).

TDP43 is crucial for several aspects of RNA processing, including RNA splicing, stability and transport (3–7). TDP43 is primarily nuclear in healthy cells, in contrast to the nuclear exclusion and cytosolic deposition of TDP43 that are characteristic of ALS (3). Although the origins of TDP43 mislocalization are unclear, deficiencies in nucleocytoplasmic transport machinery are increasingly implicated as a potential contributing factor to ALS pathology (8). Impaired nuclear import of TDP43 and other RNA binding proteins, together with dysfunction of the nuclear pore complex itself, are observed in human induced pluripotent stem cell (iPSC)-derived neurons from sporadic and familial ALS patients as well as post-mortem samples. Genetic screens in ALS model systems have repeatedly uncovered nuclear pore and nucleocytoplasmic transport factors as potent disease modifiers, confirming the significance of these pathways in disease pathogenesis (9–11).

Among the most consistent and dramatic disease modifiers to emerge from these screens is nemo-like kinase (NLK), a proline directed serine-threonine kinase that regulates cell differentiation (12), proliferation, and apoptosis (13, 14). NLK depletion mitigates disease phenotypes not just in *Drosophila* and murine models of ALS, but also murine models of spinobulbar muscular atrophy, and murine models of spinocerebellar ataxia (15–17). Conversely, NLK overexpression reduces toxicity in Huntington’s disease models, indicating context-dependent effects of NLK in neurodegeneration (18).

NLK has several predicted substrates that function in nucleocytoplasmic transport, but whether NLK itself may influence nuclear import and ALS pathogenesis remains unknown. Here, we demonstrate that NLK is a pivotal regulator of nucleocytoplasmic transport. NLK accumulation leads directly to the cytoplasmic accumulations of TDP43 and other RNA-binding proteins associated with ALS, while reduction in NLK promotes the survival of iPSC-derived neurons carrying ALS-associated mutations. Importantly, NLK is upregulated selectively in affected neurons from ALS patients and correlates with TDP43 mislocalization, implicating NLK as a key determinant of disease pathophysiology and highlighting NLK as a promising therapeutic target for ALS and related TDP43 proteinopathies.

## Results

### NLK overexpression drives mislocalization of TDP43 and other ALS-linked RNA-binding proteins

To determine if NLK is capable of directly influencing TDP43 localization, we transfected HEK293 cells with plasmids encoding FLAG-tagged wild-type (WT) NLK and examined TDP43 localization by immunofluorescence (Figure 1A). Exogenous NLK, like endogenous NLK, is both cytoplasmic and nuclear in distribution (19). As a control, we also transfected cells with plasmids encoding FLAG-tagged kinase-negative (KN) NLK, which harbors a point mutation in the active site (K155M) that abrogates kinase activity (19). NLK WT overexpression significantly decreased the nuclear-cytoplasmic (NC) ratio of endogenous TDP43 compared to NLK KN (Figure 1B). The change in TDP43 NC ratio was driven by both a reduction in nuclear TDP43 as well as a concordant increase in cytoplasmic TDP43, consistent with nuclear-cytoplasmic TDP43 mislocalization (Figure 1C-D).

**Figure 1.**
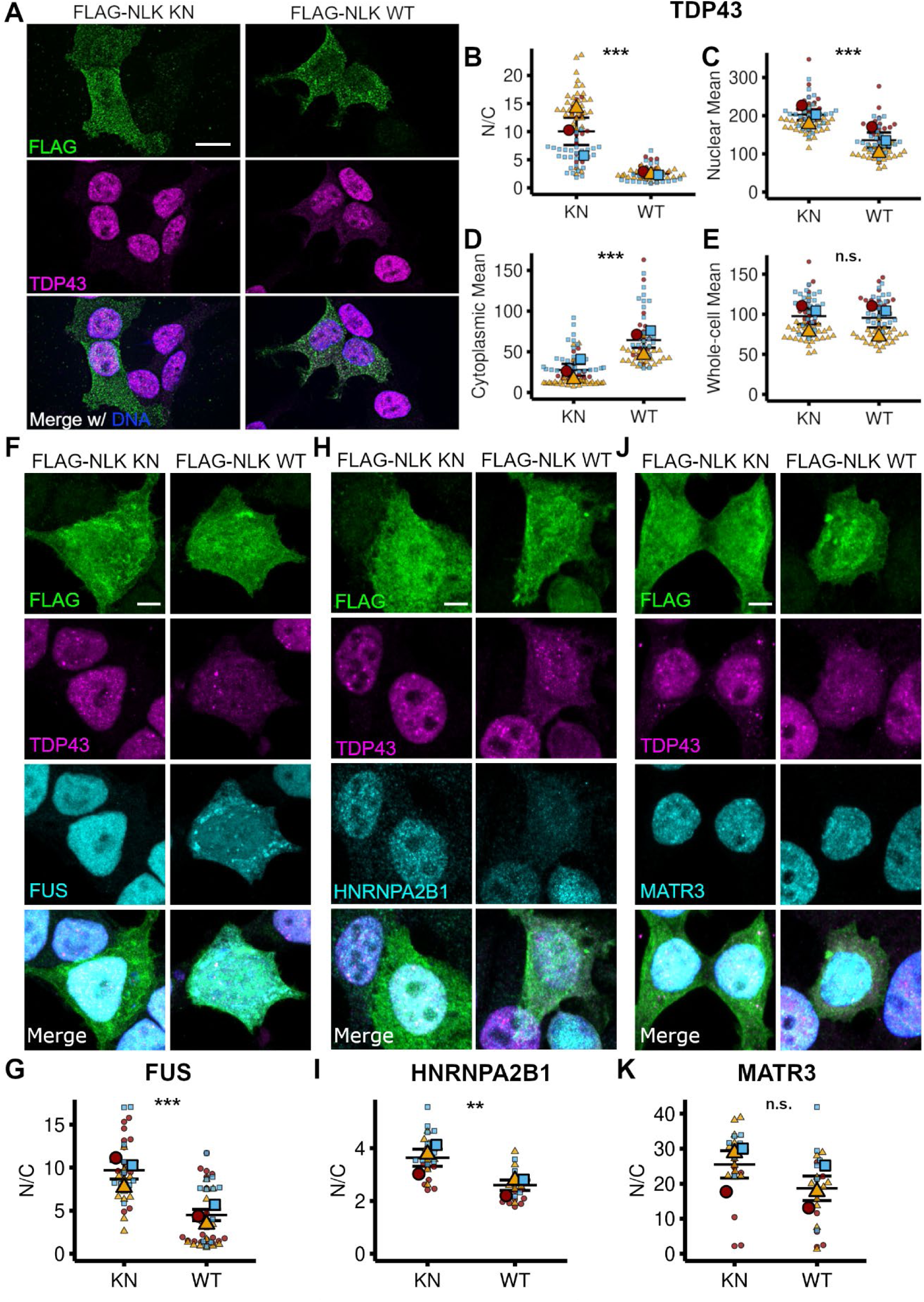
Overexpression of NLK leads to cytoplasmic accumulation of TDP43 and ALS-FTD relevant RNA-binding proteins. (**A**) HEK cells were transfected with plasmids encoding either FLAG-NLK KN (KN; kinase-negative) or FLAG-NLK WT (WT; wild-type) followed by immunofluorescence using antibodies against FLAG and TDP4343 while DNA was stained with Hoechst. Scale bar = 10μm. (**B**) Superplot of Nuclear-Cytoplasmic ratio (N/C) of TDP43 from images in (A). Line = mean, error bar = standard error. *p<0.05, ** p<0.01, ***p<0.001. (unpaired T-test with Welch’s correction). (**C**) Superplot of TDP43 nuclear intensity in cells that overexpress KN or WT NLK. Line = mean, error bar = standard error. (**D**) Superplot of TDP43 cytoplasmic intensity in cells that overexpress KN or WT NLK. Line = mean, error bar = standard error. (**E**) Superplot quantification of TDP43 whole-cell intensity in cells that overexpress KN or WT NLK. Line = mean, error bar = standard error. (**F**) HEK cells were transfected with the indicated FLAG-NLK plasmids as in (A) followed by immunofluorescence using antibodies against FLAG, TDP43, and FUS while DNA was stained with Hoechst. Scale bar = 10μm (**G**) Superplot of FUS localization in cells that overexpress KN or WT NLK. Line = mean, error bar = standard error. (**H**) HEK cells were transfected with the indicated FLAG-NLK plasmids as in (A) followed by immunofluorescence using antibodies against FLAG, TDP43, and HNRNPA2B1 while DNA was stained with Hoechst. Scale bar = 10μm. (**I**) Superplot of HNRNPA2B1 localization in cells that overexpress KN or WT NLK. Line = mean, error bar = standard error. (**J**) HEK cells were transfected with the indicated FLAG-NLK plasmids as in (A) followed by immunofluorescence using antibodies against FLAG, TDP43, and MATR3 while DNA was stained with Hoechst. Scale bar = 10μm. (**K**) HEK cells were transfected with the indicated FLAG-NLK plasmids as in (A) followed by immunofluorescence using antibodies against FLAG and MART3 while DNA was stained with Hoechst. Scale bar = 10μm. (**L**) Superplot of HNRNPA2B1 localization in cells that overexpress KN or WT NLK. Line = mean, error bar = standard error.

We next asked whether NLK WT solely affected TDP43 localization, or whether it also regulated the distribution of other ALS-associated RNA binding proteins that undergo nucleocytoplasmic transport, including HNRNPA2B1 and Matrin-3. We also examined the localization of FUS, an RNA binding protein harboring a non-classical PY-nuclear localization signal (NLS) recognized by importin-ß_2_, in contrast to the classical K/R-rich NLS in TDP43, HNRNPA2B1, and MATR3 that binds importin-ɑ (20–22). As before, HEK293 cells were transfected with KN or WT NLK, and the subcellular distribution of each protein determined by immunofluorescence (Figure 1F-K). Compared to NLK KN, NLK WT overexpression significantly reduced the NC ratio of both FUS and HNRNPA2B1 but had no significant effect on Matrin-3 localization (Figure 1F-K). Together, these findings suggest that WT NLK overexpression disrupts nuclear localization of multiple RNA-binding proteins harboring classical as well as non-classical NLSs, in a kinase-dependent manner.

### NLK overexpression disrupts nuclear import

At steady state, nuclear localization of TDP43 and other RNA-binding proteins is maintained through two competing processes: active nuclear import and passive efflux through the nuclear pore (23–25). To directly evaluate nuclear import, we co-expressed NLK together with YFP fusion proteins containing NLS sequences from TDP43 (YFP-NLS^TDP^), FUS (YFP-NLS^FUS^), Matrin-3 (YFP-NLS^MATR3^) and SV40 (YFP-NLS^SV40^, a canonical classical NLS) followed by immunofluorescence for TDP43, FUS, or Matrin-3 (Figures 2A-H). Notably, WT NLK expression drives mislocalization of YFP-NLS^MATR3^, but not endogenous Matrin-3 (Figure 1J, K), potentially due to the relatively large size of Matrin-3 (95kDa, vs 27kDa for YFP-NLS^MATR3^) (23, 24). Compared to overexpression of NLK KN, NLK WT significantly increased the NC ratio of all NLS-fusion proteins (Figure 2C). Similar to what we observed for native RNA binding proteins (Figure 1), WT NLK affects reporters containing both classical as well as non-classical NLS motifs (26). Collectively, these data indicate that WT NLK overexpression disrupts global nuclear import through its kinase activity.

**Figure 2.**
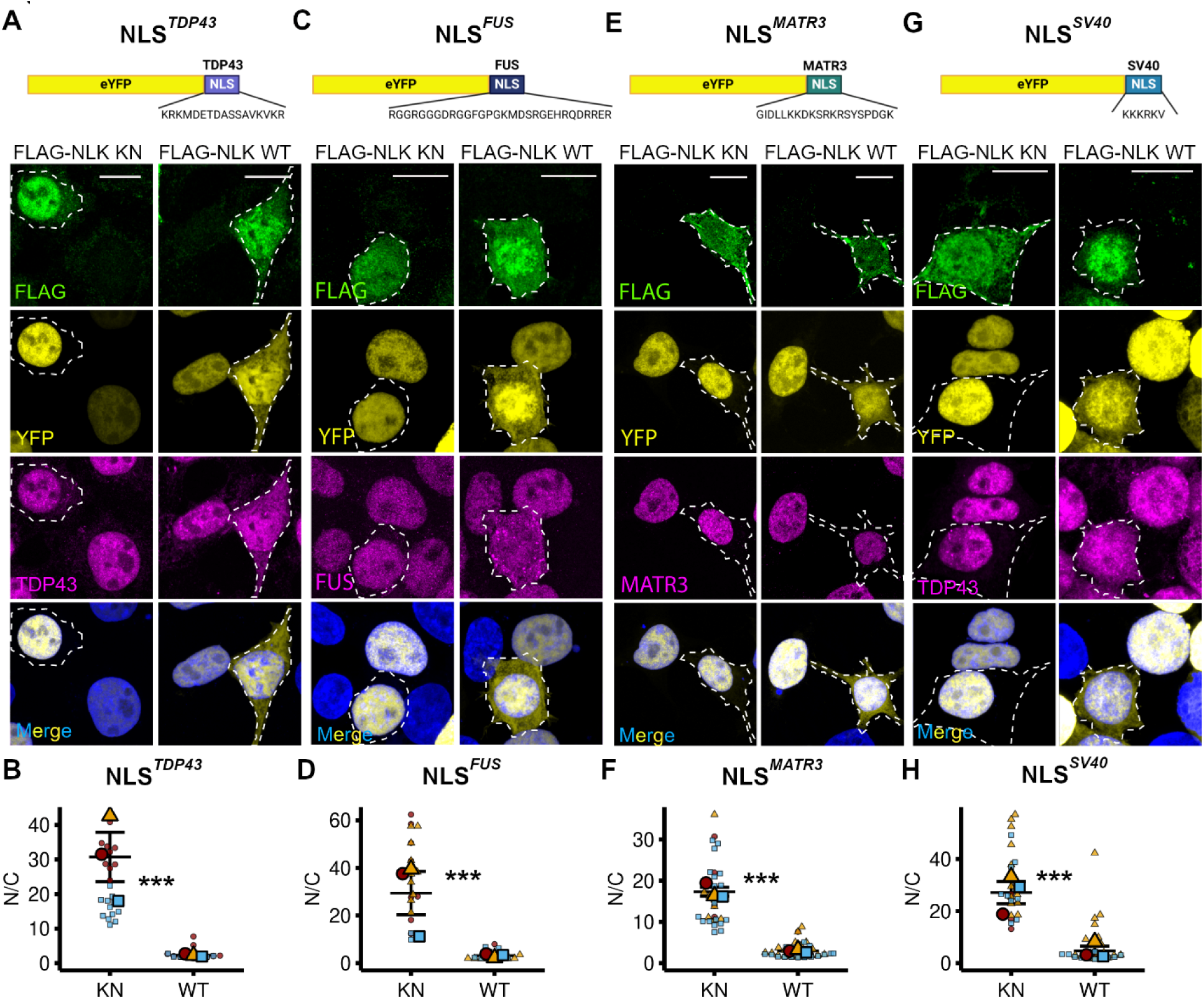
NLK overexpression disrupts NLS dependent nuclear import. (**A-D**) HEK cells were co-transfected with plasmids encoding either FLAG-NLK KN or FLAG-NLK WT and either of the following NLS-reporter eYFP-NLS^TDP43^ (A), eYFP-NLS^FUS^ (B), eYFP-NLS^MATR3^ (C), or eYFP-NLS^SV40^. Schematics created in BioRender. (D). Immunofluorescence was performed using antibodies against FLAG and either TDP43, FUS, or Matrin-3 while DNA was stained with Hoechst. Scale bar 10 uM. **(E-H**) Superplot of NLS-reporter Nuclear-Cytoplasmic ratio (N/C) from images in A-D. Line = mean, error bar= standard error. *** indicates p-value < 0.001 (unpaired t-test with Welch’s correction).

### NLK-induced TDP43 mislocalization is independent of KPNA2 nuclear accumulation

Nuclear import relies on transport receptor proteins, such as KPNA2 and KPNB1, which recognize and bind to NLS-containing proteins such as TDP43 (27). Thus, we examined the sub-cellular localization of KPNB1 and KPNA2 by immunofluorescence after overexpression of either KN or WT NLK. Overexpression of WT NLK significantly increased the NC ratio of both KPNA2 and KPNB11, driven by a significant increase in their nuclear fractions without a corresponding decrease in their cytoplasmic abundance (Figure 3A-D, S1A-B). Because of the critical importance of these factors for nuclear import, we questioned whether the mislocalization of TDP43 and other RNA binding proteins may be secondary to the observed nuclear accumulation of KPNA2 and/or KPNB1. To test this hypothesis, we took advantage of previous data showing that nuclear accumulation of KPNA2 can be reversed by the expression of the E3 ubiquitin ligase FBXW7 (28). HEK293 cells were transfected with plasmids encoding FLAG-tagged NLK WT and either mApple (negative control) or FBXW7 followed by immunofluorescence for KPNA2 (Figure 3E, F). Compared to mApple, FBXW7-V5 significantly reduced the NC ratio of KPNA2 in NLK-overexpressing cells (Figure 3G), due primarily to reduction in the nuclear KPNA2signal (Figure 3D, S1C). Despite this, FBXW7-V5 coexpression failed to significantly correct TDP43 mislocalization in cells transfected with WT NLK (Figure 3G, H, S1D). These results indicate that NLK-induced mislocalization of TDP43 does not depend on the nuclear accumulation of KPNA2.

**Figure 3.**
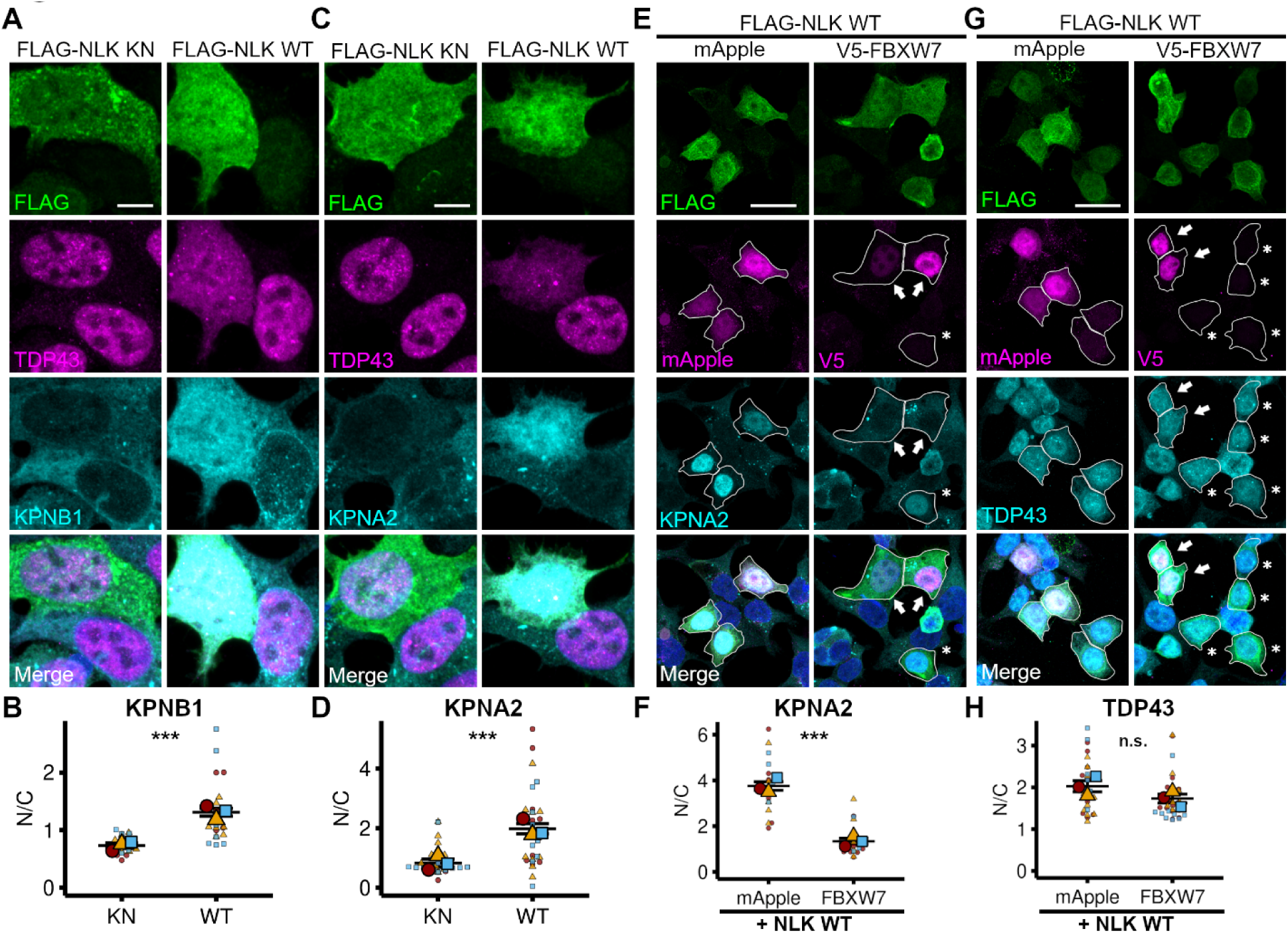
NLK-dependent mislocalization of TDP43 does not depend on nuclear accumulation of KPNA2 and KPNB1. (**A**) HEK cells were transfected with plasmids encoding either FLAG-NLK KN (KN; kinase-negative) or FLAG-NLK WT (WT; wild-type) followed by immunofluorescence using antibodies against FLAG, TDP43, and KPNA2 or KPNB1 while DNA was stained with Hoechst. Scale bar = 5μm. (**B**) Quantification of KPNA2 and KPNB1 localization in cells that overexpress NLK KN or WT. Line = mean, error bar = standard error. Statistical significance indicated as such in all sub-panels: n.s. = not significant, *p<0.05, ** p<0.01, ***p<0.001(unpaired t-test with Welch’s correction).(**C**) HEK cells were co-transfected with plasmids encoding FLAG-NLK WT and either mApple (negative-control) or V5-FBXW7 followed by immunofluorescence using antibodies against KPNA2, FLAG, and V5 while DNA was stained with Hoechst. Scale bar = 20μm. (**D**) Quantification of KPNA2 localization in cells that overexpress FLAG-NLK WT and either mApple or V5-FBXW7. Line= mean, error bar= standard error. *p<0.05, ** p<0.01, ***p<0.001 (unpaired t-test with Welch’s correction). (**E**) HEK cells were co-transfected with plasmids encoding FLAG-NLK WT and either mApple or V5-FBXW7 followed by immunofluorescence using antibodies against TDP43, FLAG, and V5 while DNA was stained with Hoechst. Scale bar = 20μm. (**F**) Quantification of TDP43 localization in cells that overexpress FLAG-NLK WT and either mApple or V5-FBXW7. Line = mean, error bar= standard error. *p<0.05, ** p<0.01, ***p<0.001 (unpaired t-test with Welch’s correction). (G) HEK cells were co-transfected as in E) followed by immunofluorescence using antibodies against KPNA2, FLAG, and V5 while DNA was stained with Hoescht. Scale bar = 20 μm.(H) HEK cells were co-transfected as in E) followed by immunofluorescence using antibodies against KPNA2, FLAG, and V5 while DNA was stained with Hoechst. Scale bar = 20μm. (I) Quantification of KPNB1 localization in cells that overexpress NLK WT and either mApple orr V5-FBXW7. Line = mean, error bar = standard error.

### NLK overexpression promotes mislocalization of Ran, Ran-GAP1, and RanBP2

Nuclear localization of receptor-bound cargo is mediated by RanGAP1 and RanBP2, nuclear pore-associated factors that regulate the Ran gradient (29)(Figure S2). Consistent with a previous screen for kinase-interacting proteins (30), both RanGAP1 and RanBP2 co-immunoprecipitated with FLAG-NLK in HEK293 cells (Figure 4A-C). NLK overexpression also promoted the accumulation of non-sumoylated RanGAP1 (Figure 4A). RanGAP1 sumoylation is critical for nuclear envelope localization and its interaction with RanBP2 (Figure 4 C, (31–33)). This prompted us to investigate the impact of NLK overexpression on the subcellular localization of RanGAP1, RanBP2, and ultimately Ran. Compared to KN NLK overexpression, WT NLK significantly disrupted the expected nuclear envelope localization of RanGAP1 and RanBP2 as measured by the ratio of nuclear rim density to cytoplasmic density (Figure 4D-G). Conversely, overexpression of NLK WT did not significantly impact FG nucleoporins as detected by MAb414 (Figure 4H, I). WT NLK overexpression also reduced the NC ratio of Ran, driven by both a decrease in nuclear Ran and an increase in cytoplasmic Ran (Figure 4J, K). As such, NLK-induced disruption of the Ran gradient correlates with disruption of the RanGAP1-RanBP2 complex and impaired nucleocytoplasmic transport.

**Figure 4.**
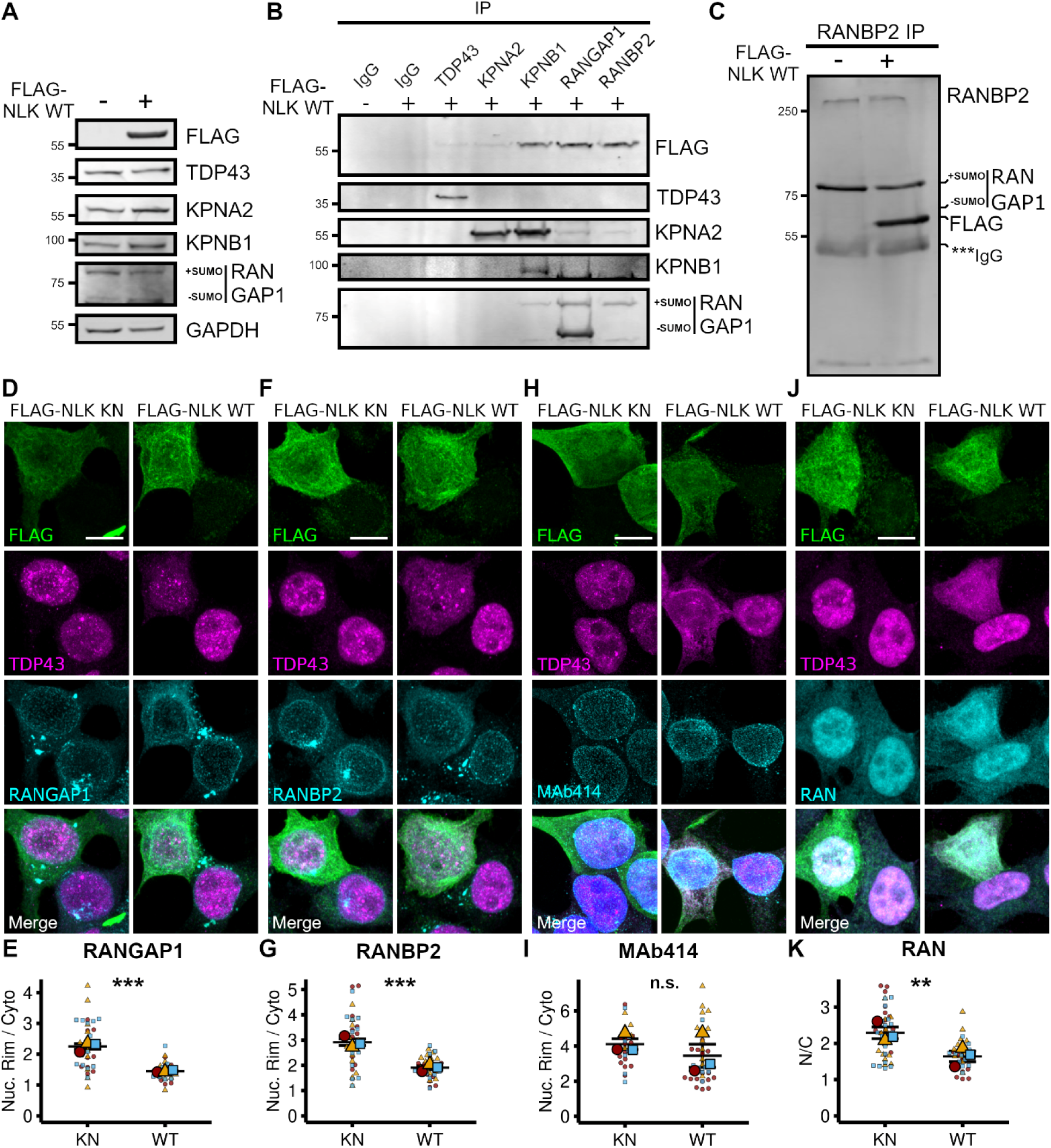
NLK overexpression mislocalizes RanBP2-RanGAP1 and disrupts the Ran gradient. (**A**) Representative western blots from HEK cells transfected with either vector (-) or FLAG-NLK WT (WT, +; wild-type). #’s to the left represent molecular weight in kDa. (**B**) Western blots after immunoprecipitation (IP) using 1μg of the indicated antibodies in 1mg of lysates from (A). (**C**) Western blots after RanBP2 co-IP from lysates in (A) were run on a 4-20% tris-glycine gel to facilitate visualization of RanBP2 and validate NLK complex formation with RanBP2-RanGAP1-SUMO. (**D**) HEK cells were transfected with plasmids encoding either FLAG-NLK KN (KN; kinase-negative) or FLAG-NLK WT (WT; wild-type) followed by immunofluorescence using antibodies against FLAG, TDP43, and either Ran, RanGAP1, RanBP2, or MAb414 (FG-nucleoporins) while DNA was stained with Hoechst. Scale bar = 10μm. (**E**) Superplot of nuclear-cytoplasmic ratio (N/C) of Ran. Line = mean, error bar= standard error. Statistical significance indicated as such in all sub-panels: n.s. = not significant, *p<0.05, ** p<0.01, ***p<0.001. *p<0.05, ** p<0.01, ***p<0.001 (unpaired t-test with Welch’s correction). (**F**) HEK cells were transfected as in (D) followed by immunofluorescence using antibodies against FLAG, TDP43, and RANBP2 while DNA was stained with Hoechst. Scale bar = 10μm. (**G**) Superplot of nuclear-rim-cytoplasm ratio (Nuc. Rim/Cyto) of RANBP2. (**H**) HEK cells were transfected as in (D) followed by immunofluorescence using antibodies against FLAG, TDP43, and MAB414 (FG-nucleoporins) while DNA was stained with Hoechst. Scale bar = 10μm. (**I**) Superplot of nuclear-rim-cytoplasm ratio (Nuc. Rim/Cyto) of MAB414. (**J**) HEK cells were transfected as in (D) followed by immunofluorescence using antibodies against FLAG, TDP43, and RAN while DNA was stained with Hoechst. Scale bar = 10μm. (**K**) Superplot quantification of nuclear-cytoplasm ratio (N/C) of RAN.

### NLK drives redistribution of mRNA and disassembles nuclear speckles

The localization of TDP43 and other RNA-binding proteins is heavily influenced not just by their NLS motifs, but also by their cognate RNA substrates (34). Based on this, we examined whether TDP43 mislocalization upon WT NLK overexpression is RNA-dependent. Initially, we transfected HEK293 cells with an EGFP-tagged variant of TDP43 harboring two mutations within RRM1 that abolish RNA binding, TDPF2L-EGFP (F147L/F149L)(35, 36), together with WT or KN NLK (Figure 5A). TDPF2L-EGFP formed phase-separated droplets that were largely restricted to the nucleus in cells co-expressing KN NLK, as in prior studies (37). However, co-transfection with WT NLK resulted in the appearance of cytosolic TDP43F2L-EGFP droplets, suggesting that TDP43 mislocalization upon WT NLK overexpression is independent of RNA binding (Figure 5A and S3A).

**Figure 5.**
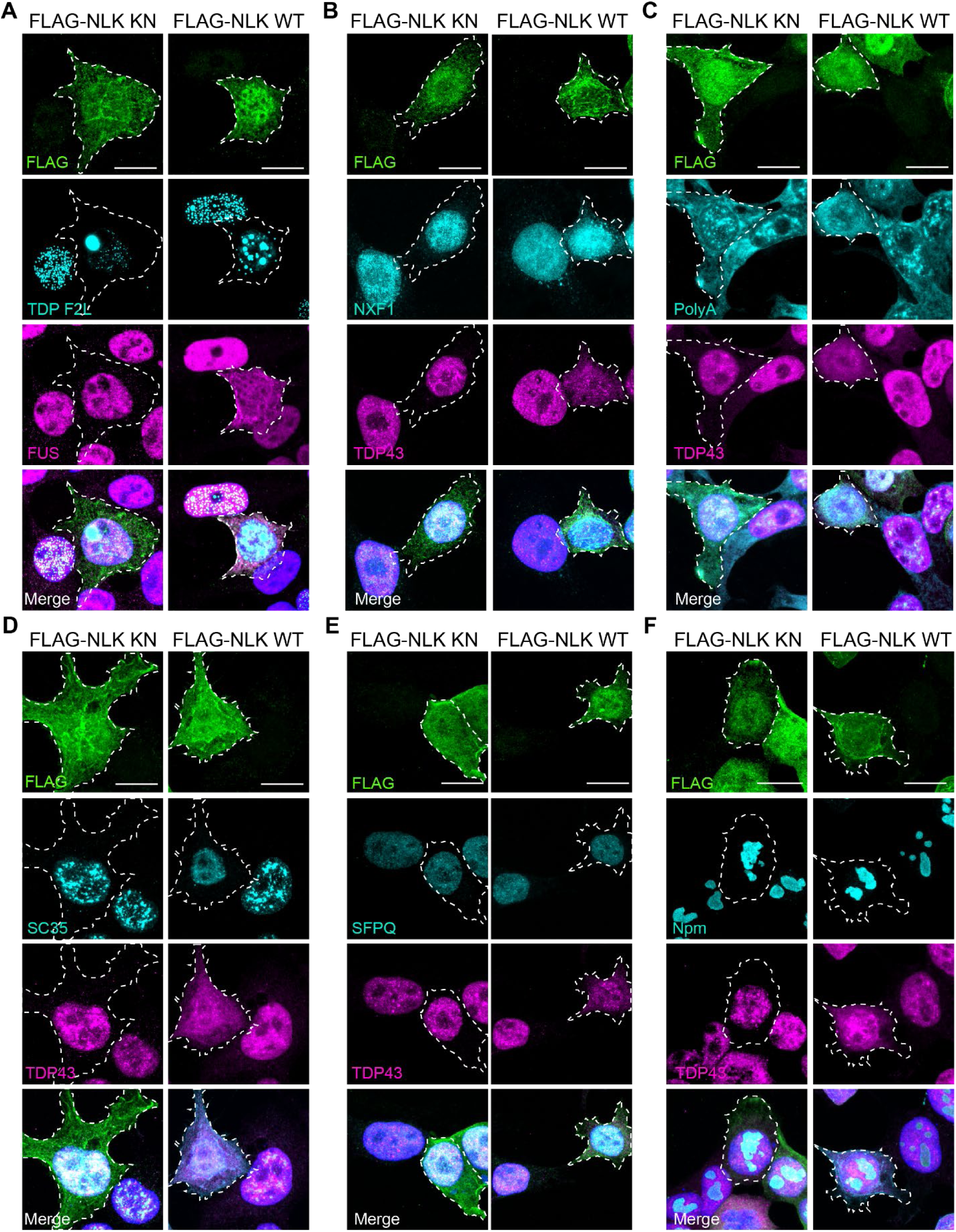
NLK overexpression drives dissolution of nuclear speckles. (**A**) HEK cells were co-transfected with plasmids encoding either FLAG-NLK KN or FLAG-NLK WT and a GFP-tagged RNA-binding mutant TDP43 (TDP43 F147/9L; F2L) followed by immunofluorescence using antibodies against FLAG and FUS and direct visualization of tagged protein while DNA was stained with Hoechst. Scale bar 10 μm. (**B**) HEK cells were transfected with plasmids encoding either FLAG-NLK KN or FLAG-NLK WT followed by immunofluorescence using antibodies against FLAG, TDP43, and NXF1. Scale bar 10 μm. (**C**) HEK cells were transfected with plasmids encoding either FLAG-NLK KN or FLAG-NLK WT followed by polyA FISH and immunofluorescence for TDP43. Scale bar 10 μm. (**D-F**) HEK cells were transfected with plasmids encoding either FLAG-NLK KN or FLAG-NLK WT followed by immunofluorescence for TDP43 and markers of speckles (SC-35) (D), paraspeckles (SFPQ) (E), or nucleolus (nucleophosmin, Npm) (F). Scale bar 10 μm.

We also investigated whether WT NLK overexpression could affect the distribution of polyadenylated (polyA) mRNA. We first immunostained for NXF1, an mRNA export factor which itself contains a PY-NLS (20) and saw that in NLK WT transfected cells, NXF1 accumulates in the cytoplasm (Figure 5B and S3B). Next, we directly assessed mRNA distribution using fluorescence in-situ hybridization (FISH). While untransfected and KN NLK-expressing cells displayed a punctate pattern of polyA mRNA within the nucleus, WT NLK overexpression resulted in a more diffuse and evenly distributed nuclear signal, and minimal changes in the polyA mRNA NC ratio (Figure 5C and S4C). Given the enrichment of polyA mRNA within nuclear speckles, and the apparent loss of such structures with WT NLK overexpression, we also immunostained WT or KN NLK-transfected cells using the SC35 antibody, which recognizes SRRM2, a core component of nuclear speckles (38). WT NLK-expressing cells, in contrast to untransfected or KN NLK-transfected cells, displayed a dramatic reduction in nuclear speckles (Figure 5D). This effect appeared to be specific for nuclear speckles, as we saw no change in other nuclear membraneless organelles such as paraspeckles (marked by SFPQ (39); Figure 5E) or nucleoli (marked by nucleophosmin (40); Figure 5F) in WT NLK-expressing cells.

### NLK overexpression disrupts nuclear import in mammalian neurons

To examine the impact of NLK overexpression on RBPs and nuclear import factors in neurons, we transfected rodent primary mixed cortical neurons with either SNAP-FLAG (SF; negative control) or SNAP-FLAG-NLK WT (SF-NLK) before immunostaining for TDP43 and other factors affected by NLK. Consistent with our results in HEK293 cells, expression of SF-NLK but not SF significantly impacted the subcellular localization of TDP43, FUS, RanGAP1, RanBP2, and Ran (Figure 6A-D). As before, the central channel of the NPC marked by FG nucleoporins was unaffected by SF-NLK expression (Figure 6E), while exogenous reporters such as YFP-NLS^SV40^ were mislocalized by SF-NLK but not SF in transfected neurons (Figure S4).

**Figure 6.**
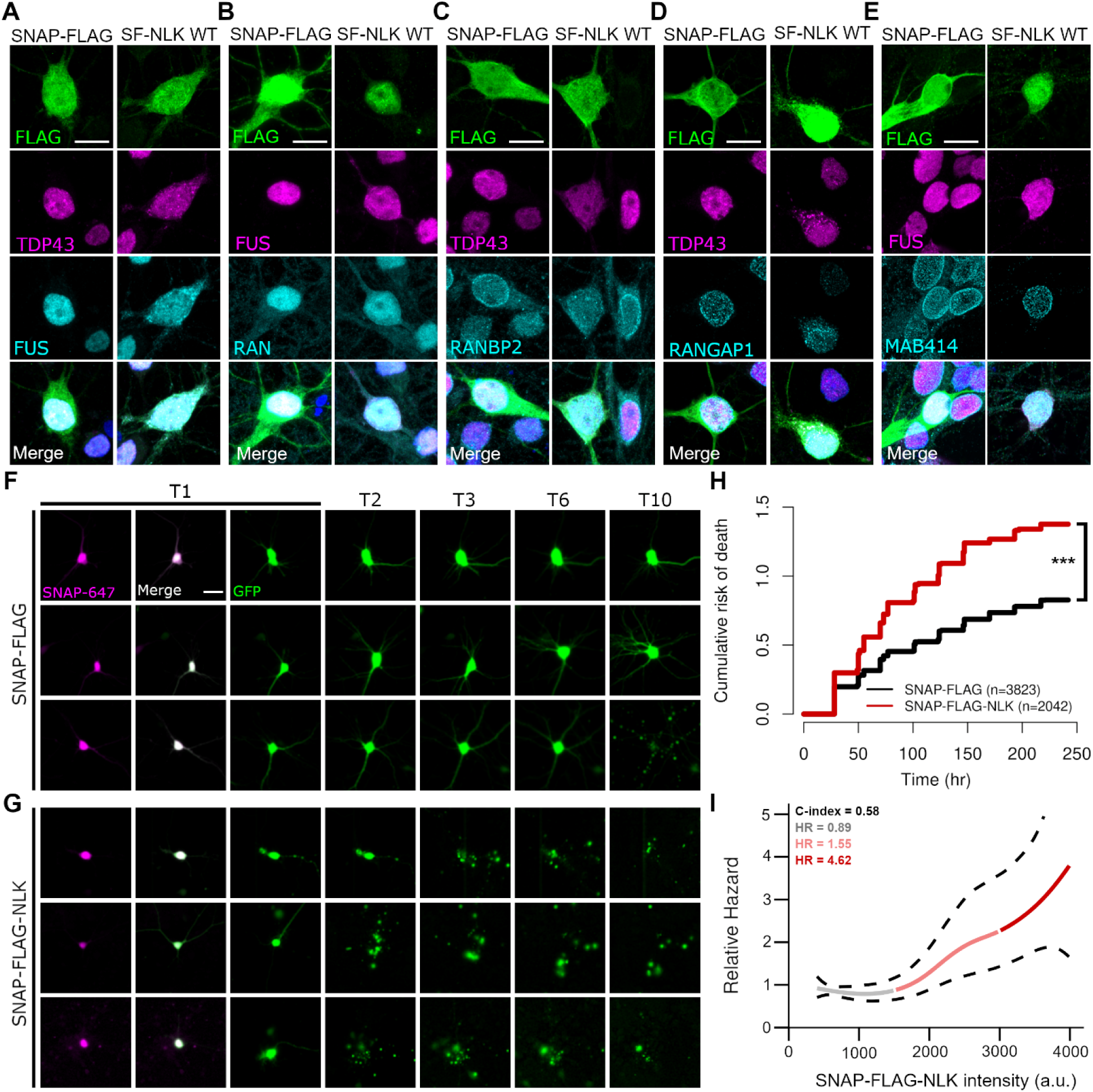
NLK overexpression disrupts nuclear import and is toxic in primary rat neurons. (**A**) Rodent primary cortical neurons were transfected with either SNAP-FLAG (SF; negative-control) or SNAP-FLAG-NLK (SF-NLK) followed by immunofluorescence using antibodies against FLAG and either TDP43, FUS, Ran, RanBP2, RanGAP1, and Mab414 while DNA was stained with Hoechst.Scale bar = 10 μm. (**B**) Rodent primary cortical neurons were co-transfected with plasmids encoding either SF or SF-NLK and EGFP (survival marker), treated with SNAP-647 dye at T1 to visualize SNAP-positive neurons, and tracked by longitudinal microscopy to determine neuron fate (see methods). Scale bar = 20μm. (**C**) Cumulative Hazard plot showing the relative risk of death in neurons that express either SF or SF-NLK. Hazard ratio = 1.616. ***p=1.494e-59. (**D**) Cox proportional hazards model predicting relative hazard associated with different intensities of SNAP-FLAG-NLK. Solid lines indicate the estimated hazard, with color gradients corresponding to intensity levels: grey (low intensity), light red (medium intensity), and red (high intensity). Dashed lines represent 95% confidence intervals for hazard estimates.

Nucleocytoplasmic trafficking is essential for maintaining protein and RNA homeostasis; thus, we predicted that NLK-induced disruption of nucleocytoplasmic transport would lead to substantial toxicity. To assess this, we utilized automated longitudinal microscopy to track hundreds of rodent primary mixed cortical neurons cultures prospectively for 10 days in culture. Neurons were transfected with SF or SF-NLK, in combination with a survival marker (GFP) enabling us to determine the time of cell death (Figure 6F, G). SF-NLK overexpression significantly increased the cumulative risk of death in transfected neurons compared to SF alone (HR=1.61, p<0.001, Cox proportional hazards analysis; Figure 6H). Since all data on survival are acquired from individual neurons, and the abundance of fluorescently-tagged proteins is directly proportional to the measured signal intensity (41), we investigated the relationship between SNAP-FLAG-NLK intensity and the hazard of the event using a Cox proportional hazards model with penalized splines (42–44). The Cox model revealed significant non-linear relationships between signal intensity and hazard, as indicated by a Likelihood Ratio Test statistic of χ² (4.06) = 25.89 (p < 0.0001), suggesting that the model fits the data well. The concordance index was 0.587, indicating a moderate ability of the model to discriminate between different hazard levels. Hazard ratios were calculated for distinct intensity segments derived from the spline model. For low signal intensities, the hazard ratio was 0.898 (95% CI: 0.378 - 2.132), indicating a slight decrease in hazard compared to the baseline. In the medium intensity range, the hazard ratio increased to 1.555 (95% CI: 0.532 - 4.545), suggesting a potential increase in risk. Notably, for high signal intensities, the hazard ratio rose significantly to 4.629 (95% CI: 0.701 - 30.584), reflecting a substantial elevation in hazard associated with increased signal intensity (Figure 6I). Collectively, our data indicate that overexpression of NLK significantly impairs the mechanisms that regulate nucleocytoplasmic transport, leading to the mislocalization of pertinent RNA-binding proteins and ultimately neuron death.

### Increased NLK levels correlate with TDP43 pathology in disease models and patients

Approximately 50% of individuals with frontotemporal lobar degeneration (FTLD) show TDP43 mislocalization as in ALS(3). In addition, up to half of people with ALS demonstrate cognitive impairment reminiscent of FTLD, while ∼1/3 of those with FTLD show motor neuron disease that is indistinguishable from ALS (3, 45). These observations, as well as shared genetics underlying both ALS and FTD, testify to the close overlap between ALS and FTLD with TDP43 pathology (FTLD-TDP) (3, 45). Therefore, to determine if NLK dysregulation may be involved in ALS/FTLD-TDP disease pathogenesis, we initially investigated *NLK* expression in *GRN* knockout mature brain organoids (mbOrgs), an FTLD-TDP model that recapitulates key pathological features of disease, including TDP43 mislocalization, phosphorylated TDP43, and characteristic missplicing of TDP43 substrate RNAs (46). Neurogenin-2 inducible cortical neurons (iNeurons) and mature cortical astrocytes (iAstrocytes) derived from isogenic wild-type (WT) or GRN-/-iPSCs were combined in fixed ratios to form mbOrgs (Figure 7A). RNA-seq revealed significantly elevated normalized counts of *NLK* in GRN-/-mbOrgs compared to WT controls (Figure 7B), a finding that was also confirmed by quantitative RT-PCR (qRT-PCR) (Figure 7C).

**Figure 7.**
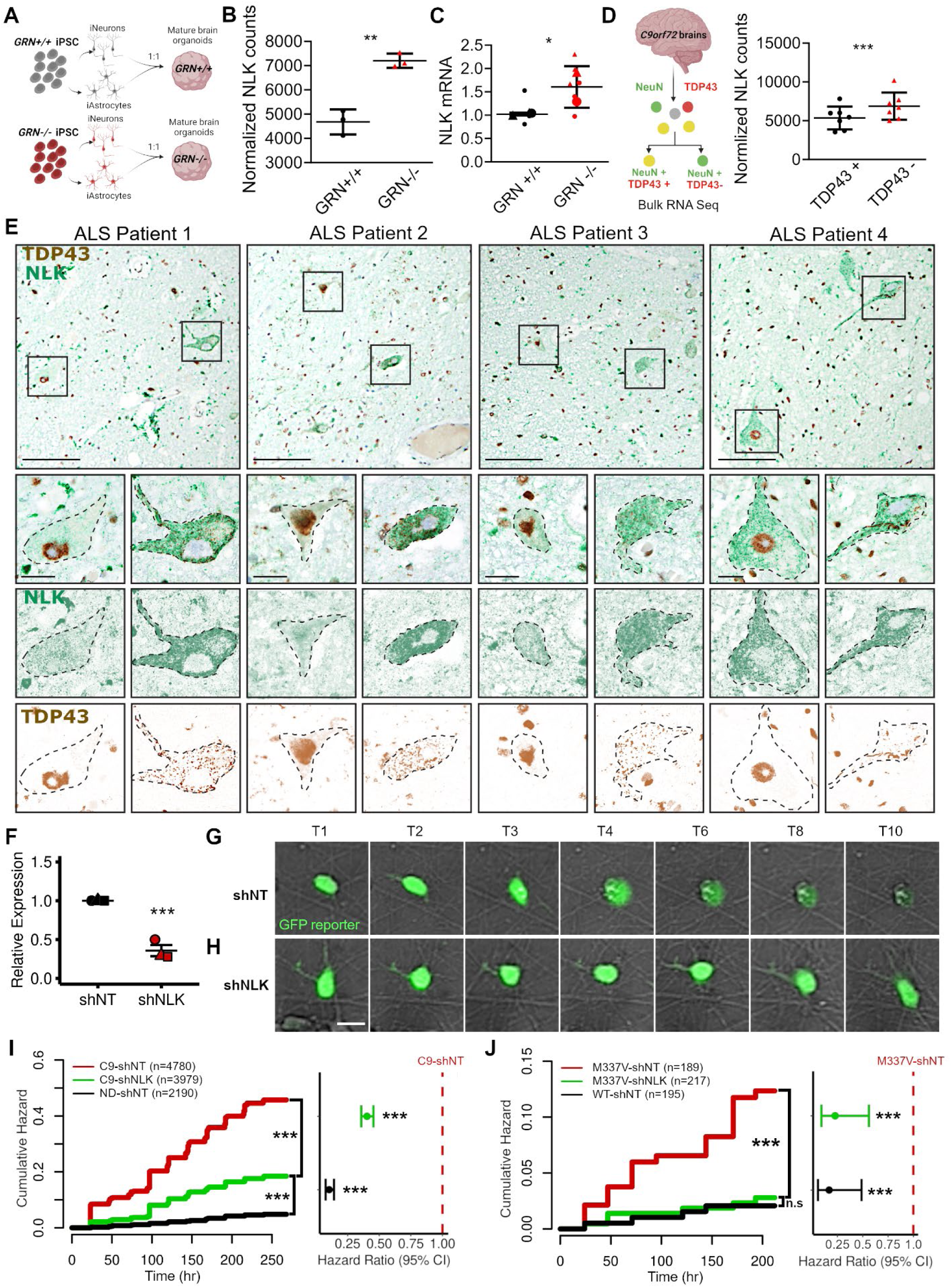
TDP43 pathology is associated with NLK overexpression in human model systems and patient samples. (**A**) Schematic of generation of mbOrgs. (**B**) Normalized NLK counts from RNASeq performed on WT or GRN^−/-^ mbOrgs. Line = mean, error bar = standard deviation. ** p< 0.01 (unpaired t-test with Welch’s correction). (**C**) QPCR was performed on WT or GRN^−/-^ mbOrgs (2 biological replicates, 3 technical replicates). Superplot of NLK levels normalized to GAPDH. Line = mean, error bar = standard deviation. * p<0.05 (unpaired t-test with Welch’s correction). (**D**) NLK normalized counts from RNAseq performed on TDP positive and negative nuclei. Line = mean, error bar = standard deviation. **** p<0.001 (Wald tests, corrected for multiple testing using the Benjamini–Hochberg method). (**E**) Dual immunohistochemistry for NLK and TDP43, performed on spinal cord tissue from four patients with sporadic ALS. Images deconvolved using FIJI. Upper panels: scale bar 100 µm. Lower panels: scale bar 20 µm. (**F**) Quantitative real-time PCR of NLK mRNA levels in iPSC derived neurons after transduction with lentivirus virus encoding either non-targeting shRNA (shNT) or NLK shRNA (shNLK). Line = mean, error bar = standard error. *** p< 0.001 (unpaired t-test with Welch’s correction). (**G**) Isogenic (WT) and mutant TDP43 (M337V) iPSC-derived neurons were transduced with virus encoding either non targeting (NT) shRNA or NLK shRNA followed by longitudinal microscopy to track neuron fate (see methods). Scale bar = 20 μm. (**H**) Cumulative Hazard plot showing the relative risk of death in ND and C9 neurons that express either shNT shRNA or shNLK (HR = 0.4, p<0.001). (**I**) Cumulative Hazard plot showing the relative risk of death in WT and M337V neurons that express either Scramble shRNA or NLKshRNA. (HR = 0.23, p=0.0013).

To further explore the link between NLK changes and TDP43 pathology, we turned to a unique dataset generated by Liu et al. in which neuronal nuclei were sorted from frontal cortices of FTLD-TDP patients into two populations — those with and without nuclear TDP43 – prior to RNA sequencing (RNA-seq) (30). Reanalysis of these data demonstrated a significant upregulation of *NLK* mRNA in nuclei lacking TDP43 (Figure 7D). Using dual-immunohistochemistry, we confirmed that spinal neurons from ALS spinal cord sections with TDP43 pathology (nuclear loss of TDP43 with cytosolic inclusions) exhibited more intense staining for NLK in comparison to unaffected neurons present in the same section (Figure 7E and S5C). Dual staining for NLK and TDP43 was also performed in sections from four control patients without spinal pathology (Figure S5D5), showing no clear differences in NLK abundance. These results demonstrate a clear relationship between elevated NLK, at both the mRNA and protein levels, and TDP43 mislocalization in FTLD-TDP and ALS.

Together, these data show that NLK is upregulated in human patients and in disease models featuring TDP43 pathology, in accord with our data demonstrating TDP43 mislocalization upon NLK overexpression.

### NLK reduction improves survival in iPSC-derived neuron models of ALS

Given the detrimental effects of NLK overexpression on nucleocytoplasmic transport (Figures 1-6) and neuron survival (Figure 6), and the elevated *NLK* mRNA and protein observed in patients and disease models (Figure 7A-E), we asked whether targeting *NLK* could ameliorate disease phenotypes in ALS/FTLD-TDP models. First, we examined the survival of iPSC-derived neurons carrying *C9ORF72* expansions, the most prevalent mutation underlying familial ALS and FTLD-TDP in Europe and North America (47, 48). Non-disease and *C9ORF72* mutant neurons were transduced with virus encoding non-targeting or *NLK* shRNA, resulting in a ∼50% reduction in NLK RNA levels compared to non-targeting shRNA (Figure 7F). Individual neurons were followed by automated microscopy for 10 days, as before (Figure 7G), and differences in survival assessed via Cox proportional hazards analysis. Three separate lines of *C9ORF72* mutant neurons exhibited significantly higher cumulative risks of death compared to unrelated non-disease neurons (Figure 7H and S6A-C). Transduction with *NLK* shRNA significantly reduced the cumulative risk of death in all three lines of *C9ORF72* neurons (HR= 0.403, p= 7.71e-50). To examine if NLK reduction is neuroprotective outside of *C9ORF72* mutations, we utilized an isogenic pair of *TARDBP* mutant iPSC-derived neurons that were created by CRISPR/Cas9 genome engineering (49). In comparison to isogenic controls (“WT”), *TARDBP* mutant (M337V) neurons transduced with lentivirus expressing non-targeting shRNA exhibited a significantly higher cumulative risk of death (Figure 7I). As with *C9ORF72* mutant neurons, however, transduction with *NLK* shRNA-expressing virus significantly extended the survival of M337V neurons (HR = 0.227, p= 0.0013). Collectively, these data imply that *NLK* overexpression is toxic, while *NLK* reduction promotes neuronal survival in models of ALS and FTLD-TDP.

## Discussion

Our findings indicate that NLK overexpression disrupts the nuclear import and localization of TDP43 and related RNA-binding proteins, including FUS and HNRNPA2B1, in a kinase-dependent manner. These effects correlate with the mislocalization of the RanBP2-RanGAP1 complex and collapse of the Ran gradient, both of which are crucial for functional nucleocytoplasmic transport. As expected based on these observations, NLK dose-dependently increased the risk of death in primary rodent neurons. Notably, we uncovered elevated NLK expression selectively in neurons with TDP-43 pathology both at the RNA level (in FTLD-TDP frontal cortex) and at the protein level (in ALS spinal cord). NLK was also upregulated in a brain organoid model of FTLD-TDP, while reducing NLK in human iPSC-derived neurons carrying disease-associated mutations prolonged survival. These results suggest that NLK may contribute to the pathology of ALS-FTLD-TDP by impairing nucleocytoplasmic transport and promoting neurotoxicity. Targeting NLK protein levels or kinase activity could thus represent a novel therapeutic approach for ALS, FTLD-TDP and other TDP43 proteinopathies.

Disruption of the Ran gradient, mislocalization of nuclear pore components, and disturbance of nuclear pore architecture — all of which we observed upon NLK overexpression — have likewise been described in ALS/FTLD-TDP samples and disease models (9–11, 50). There is no predicted NLK phosphorylation site within TDP43 itself; rather, we suspect that NLK may interfere with the nucleocytoplasmic transport of several RNA binding proteins and other factors by acting on integral components of the nuclear pore. We and others noted a direct interaction between NLK and the RanGAP1-RanBP2 complex ((30); see Figure 4), which is critical for maintaining the Ran gradient and nuclear import/export (51). NLK overexpression prevented RanGAP1 sumoylation (Figure 4A) and reduced the amount of RanGAP1 in complex with RanBP2 (Figure 4C). In the absence of sumoylation, RanGAP1 is unable to bind RanBP2 or localize to the nuclear envelope (Figure 4C, (31–33). One possibility is that NLK directly phosphorylates RanGAP1, inhibiting its GAP activity (50) as well as its sumoylation, thereby disrupting its interaction with RanBP2 at the nuclear pore complex. Alternatively, NLK may negatively regulate Ubc9, a SUMO E3 ligase required for recruitment of RanGAP1 to the RanBP2 complex. None of these possibilities are mutually exclusive; NLK may impact nucleocytoplasmic transport through more than one overlapping mechanism.

At baseline, NLK is highly expressed in the CNS, and its expression is unregulated by oxidative and osmotic stress (52, 53), conditions associated with the cytosolic accumulation of TDP43 and other, predominantly nuclear, RNA binding proteins. NLK interacts with several proteins associated with neurodegenerative diseases, including poly-Q expanded androgen receptor in spinobulbar muscular atrophy (SBMA) and ataxin-1 in spinocerebellar ataxia 1 (SCA1) (16, 17); accumulations of these proteins may also contribute to changes in NLK expression and/or activity. Although previous studies have failed to detect NLK upregulation in disease, in most cases these investigations are limited to evaluation of NLK levels (RNA or protein) in bulk tissue. In contrast, our work revealed NLK upregulation solely within affected neurons displaying TDP43 pathology, in association with evidence of disrupted nucleocytoplasmic trafficking in the same cells, suggesting that NLK expression changes are restricted to cells with TDP43 redistribution. Outside of changes in expression, NLK may also be inappropriately activated in disease via diverse stimuli. Proinflammatory factors such as transforming growth factor ß (TGFß), interleukin-6 (IL6) and Wnt all activate NLK through MAPK-dependent signaling (54). Notably, TGF-ß is upregulated in ALS (55), and aberrant Wnt signaling is associated with TDP43 mislocalization in cellular and animal models (56).

NLK is an essential kinase — homozygous *Nlk* deletion results in death *in utero* or postnatally, depending on the mouse strain (57, 58). Conditional *Nlk* knockout in adult animals is well-tolerated, however (59), and *Nlk* haploinsufficiency (*Nlk*+/-) is protective in several neurodegenerative disease models, including TDP43-overexpressing mice (15–18). Similarly, partial *NLK* knockdown in our investigations was associated with extended survival in *C9ORF72* and *TARDBP* mutant human iPSC-derived neurons and was safe in control neurons. These data indicate that even modest reductions in *NLK* may be sufficient for mitigating neurodegeneration in ALS and FTLD-TDP.

Although prior interaction studies highlighted several potential partners for NLK, its substrates in neurons and phosphorylation sites within these substrates remain largely unexplored. RanBP2 and RanGAP1 are two potential targets for NLK with clear connections to nucleocytoplasmic transport, but NLK is also likely to act on distinct substrates and disease-related pathways, including the integrated stress response and autophagy (15). One advantage of therapeutic strategies that act on upstream signaling factors such as NLK is the capacity to influence several neuroprotective mechanisms simultaneously. Nevertheless, not all downstream events are likely to be beneficial(18), emphasizing the need for further investigations into NLK targets and the impact of NLK-mediated phosphorylation on their function and contribution to disease.

## Methods

### Sex as a biologic variable

Our study examined tissue from both male and female patients, and similar findings are reported for both sexes.

### HEK293 cell culture and transfection

Human embryonic kidney (HEK) 293T were cultured in DMEM (Gibco, 11995065) supplemented with 10% FBS (Gibco, ILT10082147) at 37°C in 5% CO2. Cells were transfected with Lipofectamine 2000 (Invitrogen, 11668027) according to the manufacturer’s instructions. For experiments with YFP-NLS reporters, the lipofectamine solution was divided toto ensure there would be both cells co-transfected with NLK and reporter as well as cells transfected with the reporter alone. Of the lipofectamine solution, 75% contained two plasmids (NLK and reporter) while the remaining 25% contained reporter alone. The lipofectamine solution was then combined and briefly mixed before adding directly to cells. After 24 hours, transfected cells were split in media containing poly D-lysine (1:500, cat Millipore A-008-E) onto coverslips coated with laminin (1:100, Sigma L2020-1MG). Cells were cultured an additional 24 hours then immunocytochemistry was performed as detailed below.

### Primary neuron cell culture and transfection

Cortices from embryonic day (E)19-20 Long-Evans rat embryos were dissected and disassociated, and primary neurons were plated at a density of 6×10^5^ cells/mL in 96-well plates or in 24 well plate on coverslips. At *in vitro* day (DIV) 4, neurons were transfected using Lipofectamine 2000 as previously described (49, 60). For experiments with the YFP-NLS reporter, the transfection mix was split as described above for HEK cells. Following transfection, cells were placed in Neurobasal Complete Media (Neurobasal (Gibco 21103-049), 1x B27 Supplement (Gibco, 17504-044), 1x Glutamax, 100 units/mL Pen Strep). Cells were fixed for immunocytochemistry 48 hours after transfection.

### iPSC maintenance and differentiation

Creating Neural Progenitors: iPSCs were dissociated using Accutase, counted, and plated at 32,000 cells/mL in E8 media with ROCK inhibitor. Differentiation was then induced with doxycycline for 48 hours (2 μg/mL; Sigma, #D3447). Differentiating neural progenitors (NP) were dissociated using Accutase after two days of doxycycline induction and frozen for future experiments.

Differentiating Neural Progenitors: Previously frozen neural progenitors were thawed in E8 media with ROCK inhibitor and incubated for 24 hours. Subsequently, the media was changed to N2 media containing 1x N2 Supplement (Gibco, #17502-048), 1x NEAA Supplement (Gibco, #11140-050), 10 ng/mL BDNF (Peprotech, #450-02), 10 ng/mL NT3 (Peprotech, #450-03), 0.2 μg/mL Laminin (Sigma, #L2020), and 2 μg/mL doxycycline (Sigma, #D3447). On Day 2, the media was replaced with transition media (1:1 full E8 media and DMEM/F12 (Gibco, #11320-033)). On Day 3, the media was switched to B27 media, prepared with 1x B27 Supplement (Gibco, #17504-044), 1x Glutamax Supplement (Gibco, #35050-061), 10 ng/mL BDNF, 10 ng/mL NT3, 0.2 μg/mL Laminin, 1x CultureOne (Gibco, #A33202-01), and Neurobasal-A (Gibco, #12349-015). On Day 6, cells were transduced with the virus (University of Michigan Vector Core) and maintained in the same media for the remainder of the experiment.

### iPSC cell lines

The cell lines 1021 and 793 (controls) and 883 and 321 (C9ORF) were generated and characterized as previously described (7).. The CS52i and corrected cell lines were obtained from Cedars Sinai. See table below for additional details.

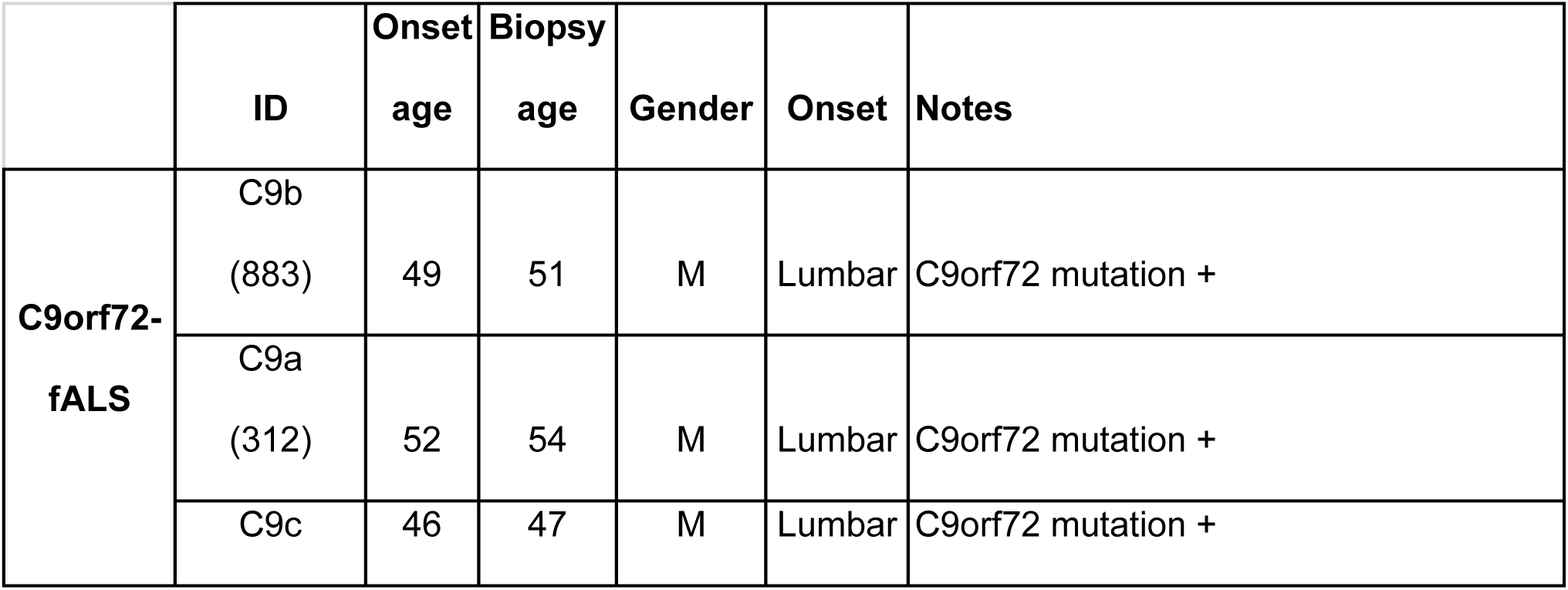
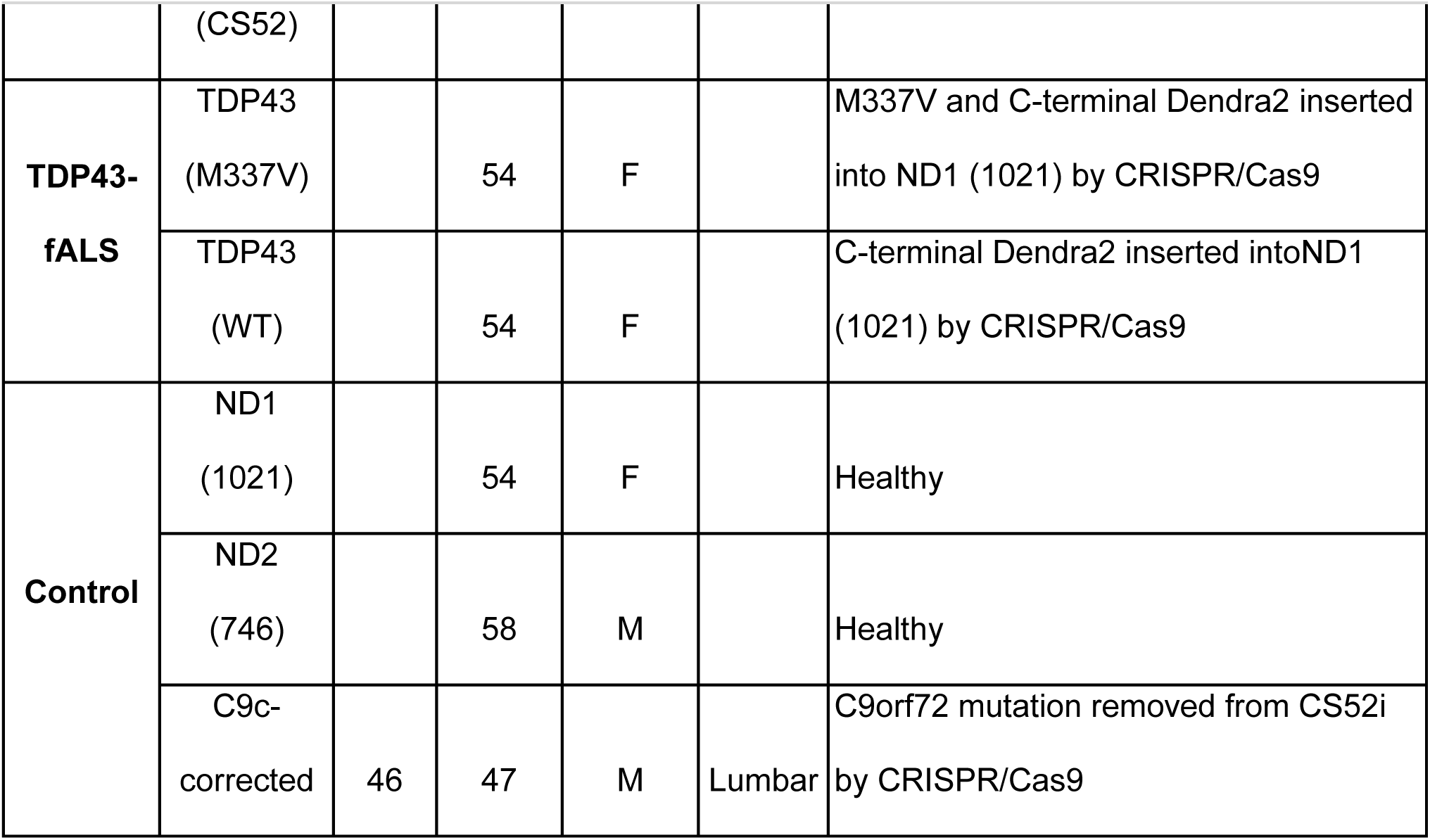

### mbOrg generation

Isogenic human iPSC line WTC11 (GRN+/+ and GRN−/−) were generated by Dr. Bruce R. Conklin, as previously described (61). iPSCs were cultured and maintained in StemMACS™ iPS-Brew XF (Miltenyi Biotec, 130-104-368) media on 6-well cell culture plates (GenClone, 25-105MP) coated with Vitronectin (Gibco, A14700) in DPBS. iPSCs were dissociated and passaged using EDTA (Invitrogen, AM9260G) in DPBS.

iA and iN were differentiated separately, plated in a 1:1 ratio, and aged for 4 weeks as previously described (46). For iN differentiation, iPSCs were transduced with NGN2-expressing lentivirus constructs and a previously published protocol was followed for differentiation (61). Briefly, iPSCs are expanded, dissociated, and replated on Matrigel-coated plates. Cells are grown in specialized iNeuron induction media (DMEM-F12 + Glutamax; Gibco, 10565-018), N-2 supplement (Gibco, 17502-048), MEM-NEAA (Gibco, 11140-050) containing doxycycline (Sigma, D3072) for 72 h, with media changed every 24 h. Cells are then dissociated using Accutase and plated together with iA. Cortical-like iA are generated as previously described (62). Human induced pluripotent stem cells (GRN+/+ and GRN-/-) were grown on vitronectin coated tissue culture plates using StemMACS™ iPS-Brew XF (Miltenyi Biotec, 130-104-368) media. On day 0 of differentiation, iPSCs are dissociated into small aggregates and transferred in neurosphere induction medium (NSIM) (DMEM-F12/Neurobasal-A at 1:1) plus SMAD inhibitors SB431542 and DMH1. On day 7, spheroids are transferred to Matrigel-coated tissue culture plates with NIM. On day 14, rosette clusters were mechanically removed and transferred to tissue culture flasks with NIM plus FGFb (Peprotech, 100-18B). Media was changed every 72 hours. On day 20, spheroids were triturated into a single cell suspension and transferred to a new untreated cell culture flask with astrocyte media (ASM) [(DMEM-F12 (Gibco, 10565-018), N2 Supplement (Gibco, 17502-048), B27 -Vit.A Supplement (Gibco, 12587-010), Heparin (Stemcell Tech, 07980)) plus Y27632 (Tocris, 1254)]. From Day 28 to 180, spheroid aggregates were maintained in suspension with ASM plus EGF and FGFb (Peprotech, 100-15 & 100-18B) with media changes every 4-5 days. Spheroid aggregates were triturated every 7-10 days and transferred to new untreated tissue culture flasks. Spheroids are triturated every 7– 10 days and transferred to new tissue culture flasks. In order to generate mbOrgs, iN and iA are plated together at a 1:1 ratio on Matrigel-coated 24-well plates at a collective density of ∼1×106 cells/well. Cells are maintained in BrainPhys Complete medium with a 50% medium change every 72 h. mbOrgs will be created on 384-microwell plates as described previously (46), resulting in mbOrgs that are highly uniform in size (∼500 m) after brief spin-down and 1–2 days of subsequent culture. These 3D co-cultures were aged for 4 weeks.

### Longitudinal microscopy

At in vitro day (DIV) 4, neurons were transfected with 100ng of a control fluorescent plasmid to mark cell bodies and 100ng of an experimental construct using Lipofectamine 2000 as previously described (43, 49, 62). Neurons were imaged as described previously (43, 49, 60, 63) using a Nikon Eclipse Ti inverted microscope with PerfectFocus3a 20X objective lens and either an Andor iXon3 897 EMCCD camera or Andor Zyla 4.2 (+)sCMOS camera. A Lambda CL Xenon lamp (Sutter) with 5 mm liquid light guide (Sutter)was used to illuminate samples, and custom scripts written in Beanshell for use in micromanager controlled all stage movements, shutters, and filters. For automated analyses of primary neuron survival, custom ImageJ/FIJI macros and Python scripts were used to identify neurons and draw cellular regions of interest (ROIs) based upon size, morphology, and fluorescence intensity. Custom Python scripts were used to track ROIs over time, and cell death marked a set of criteria that include rounding of the soma, loss of fluorescence and degeneration of neuritic processes (63, 64). For iNeuron survival, brightfield and fluorescent images for survival experiments started on Day 14 and continued for 10 days. For manual analysis of iNeuron survival, image time-series were processed by flat-field correction and image registration, followed by programmatic de identification for blinded analysis. Time-series images were uploaded to a browser-based server where a trained user manually counted survival using the point tracking mode. GFP-positive cells were identified at T1, and cell fate was tracked in brightfield images, where neuron death (uncensored event) was recorded at the appropriate time point. Living neurons at the final time point were considered right-censored. Cox proportional hazards model was applied to the datasets, stratifying by biological replicates, and the corresponding hazard plots were generated in R.

### Immunocytochemistry

Cells were fixed with 4% paraformaldehyde (PFA; Sigma, P6148) for 10m, rinsed with PBS, and permeabilized with 0.1% Triton X-100 (Bio-rad, 161-0407) for 10m. Cells were then blocked in 3% bovine serum albumin (BSA; Fisher, BP9703-100) in PBS at RT for 1h before incubation O/N at 4°C in primary antibody diluted in 3% BSA (See table for additional details). Cells were then washed 3 times in PBS and incubated at RT with secondary antibodies diluted 1:250 in 3% BSA for 1h. Following 3 washes in PBS containing 1:10,000 Hoechst 33258 dye (Invitrogen, H3569), cells were mounted on Superfrost Plus Microscope Slides (Fisher, 1255015) with Prolong Gold Antifade Mounting Reagent (Fisher, P10144) and imaged as described below. Images were analyzed by collecting measurements within cellular regions of interest using custom ImageJ/FIJI macros. Graphs were made using R.

### Fluorescence in situ hybridization

Cells were fixed with 4% paraformaldehyde (Cat #) for 10 m, rinsed with PBS and permeabilized with 0.1% Triton X-100 (Bio-rad, 161-0407) for 10m. Cells were then washed with 2X SSC (2 x 5 m) and incubated in hybridization buffer for 2 hours at 37 degrees (8.75% dextran, 1.75X SSC, 17.5% formamide, 0.5 ug/ul yeast tRNA, 10 mmol RVC, 0.1% BSA, 0.002 ug/ul Cy3-oligo (dT) 30 probe). Cells were then washed with successive SSC rinses (4X SSC 15 m, 2X SSC 15 m, 2X SSC 15m). Then immunocytochemistry was performed as above, beginning with blocking in 3% BSA. All solutions prior to hybridization were DEPC treated.

### Immunohistochemistry

Duplex immunohistochemistry was performed on a Ventana Discovery Ultra stainer (Indianapolis, IN). Slides were dewaxed, rehydrated and subjected to heat induced epitope retrieval on board the stainer. Slides were then subjected to sequential incubation with NLK (Rabbit polyclonal antibody, AbCam, Cambridge, MA, Ab26050, 1:250, 32 minutes) and polymer goat anti-rabbit IgG conjugated to HRP (Ventana) and developed with Discovery Green chromogen (Ventana). After an additional round of heat induced epitope retrieval to remove the NLK primary antibody-secondary antibody complex, the slides were stained with TDP43 (Rabbit polyclonal antibody, Proteintech, Rosemont, IL, 10782-2-AP, 1:2000, 20 minutes), polymer goat anti-rabbit IgG conjugated to HRP and developed with Discovery Brown chromogen (Ventana). Slides were then countered stained with hematoxylin and coverslipped.

### RNA isolation for bulk RNAseq and RT-PCR

RNA extraction was performed using the Trizol/phenol-Chloroform method (Sigma, T9424) as previously described (62) and according to manufacturer specifications. Each sample contained ∼60 mbOrgs, totaling ∼3 ×10^6^ of cells per sample. The extracted RNA was used as a template for the synthesis of complementary DNA (cDNA) through reverse transcription, using iScript^TM^ cDNA Synthesis Kit (Bio-Rad Cat#1708891) according to the manufacturer’s protocol.

### Quantitative PCR

cDNA samples were treated for genomic DNA contamination using DNA-*free*™ DNA Removal Kit (Invitrogen Cat#AM1906) per manufacturer’s instructions. The cDNA was then diluted to a concentration of 5 ng/µl and 4 µl of each sample (total of 20 ng) were aliquoted in a MicroAmp™ Optical 96-Well Reaction Plate (Thermo Scientific Cat#N8010560) in technical duplicates. Samples were processed using 2X SYBR Green qPCR Master Mix Assay and quantitative PCR was run on QuantStudio 6 Real-Time PCR system following manufacturer’s instructions. Data analysis was carried out applying the Pfaffl mathematical model for relative transcript quantification (65) using GAPDH as a housekeeping gene.

### RNA Sequencing

RNA sequencing and analysis of RNA integrity was analyzed on an Agilent 2100 Bioanalyzer using RNA 6000 Nano kit (Agilent, 5067-1511). Only samples with an RNA integrity number (RIN) ≥ 9.4 were used to perform bulk RNA sequencing. Nugen Universal Plus (Tecan) was used as a library kit and libraries were sequenced on a SP300 flow cell of the Illumina NovaSeq 6000 machine with a paired end 150 bp sequencing strategy (average depth 90 million reads/sample) at UCSF Genomics Core Facility. Genome was aligned to Ensembl Human.GRCh38.103. Kallisto 0.46.01 was used to generate transcript abundance files for each sample. Transcript counts files for each sample were generated using txImport and transcript differential analysis was performed using DESeq2 v1.24.0. A total of 6 samples were spread across two conditions.

### Confocal microscopy

Confocal images were taken on a Nikon AXR NSPARC confocal system with a 60x NA1.42 Oil/DIS PLan-Apochromat Lambda D objective with a working distance of 1.5 mm, and a 40x CFI Apochromat LWD Lambda S objective with a working distance of 0.30 mm.

### Light microscopy

Whole-slide images were generated by the University of Michigan Digital Pathology group within the Department of Pathology using an Aperio AT2 scanner (Leica Biosystems) equipped with a 20x NA 0.75 Plan Apochromat objective. 40x scanning is achieved using a 2x optical magnification changer. Resolution is 0.25 μm/pixel for 40x scans. Focus during the scan was maintained using a triangulated focus map built from individual focus points determined in a separate step before scanning was started. Proprietary software was used for image processing during the acquisition. TIFs were taken of the entire ventral horn of all cases, then a pathologist selected each neuron, manually drew an ROI, and annotated the TDP status (pathology or no pathology) using custom Fiji scripts. Then deconvolution and analysis were performed using custom Fiji scripts.

High quality images for figures were acquired on an Olympus BX51 light microscope equipped with a UPlanSApo100x oil objective with a numerical aperture of 1.40 and a working distance of 0.12 mm. Image deconvolution was performed using Fiji.

### Antibodies

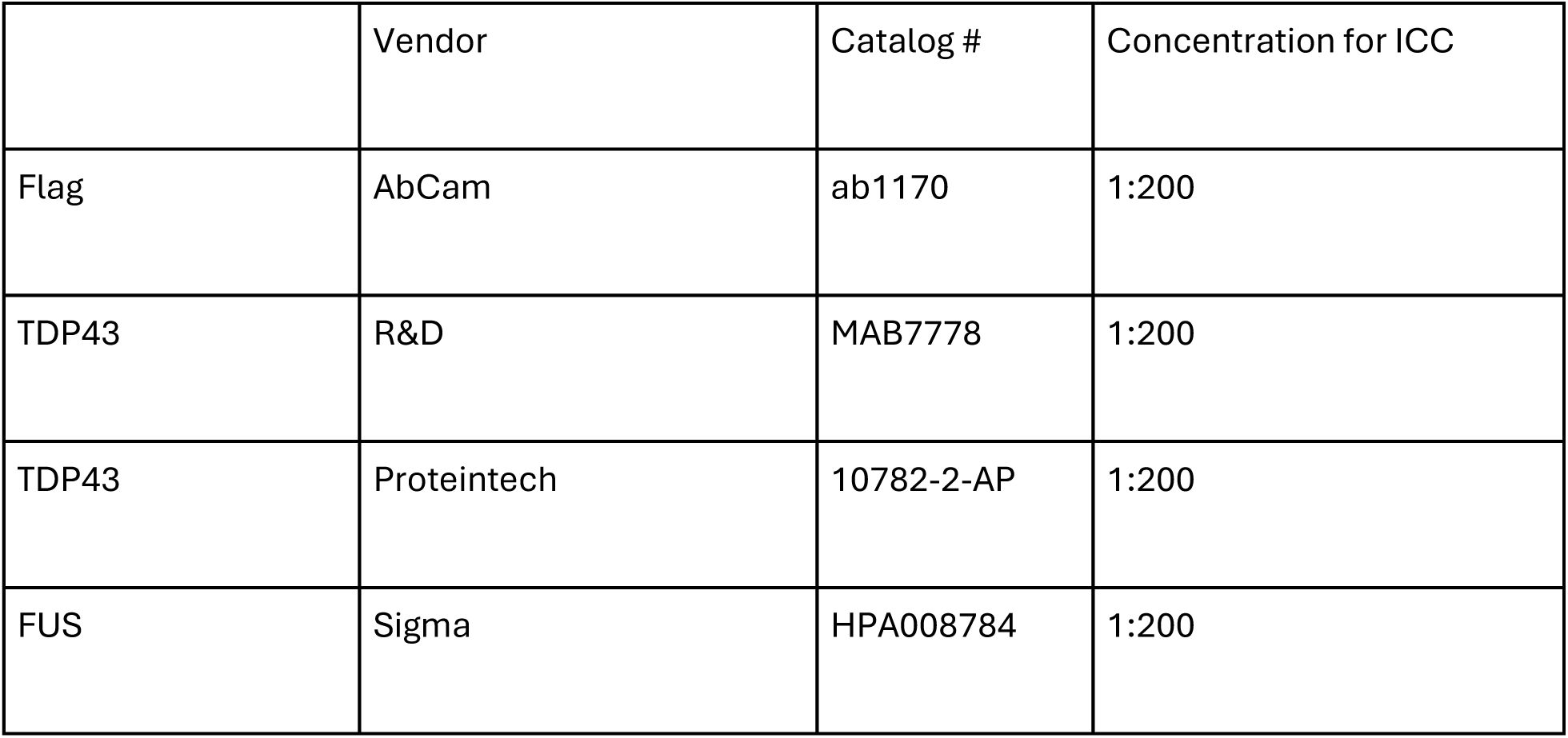
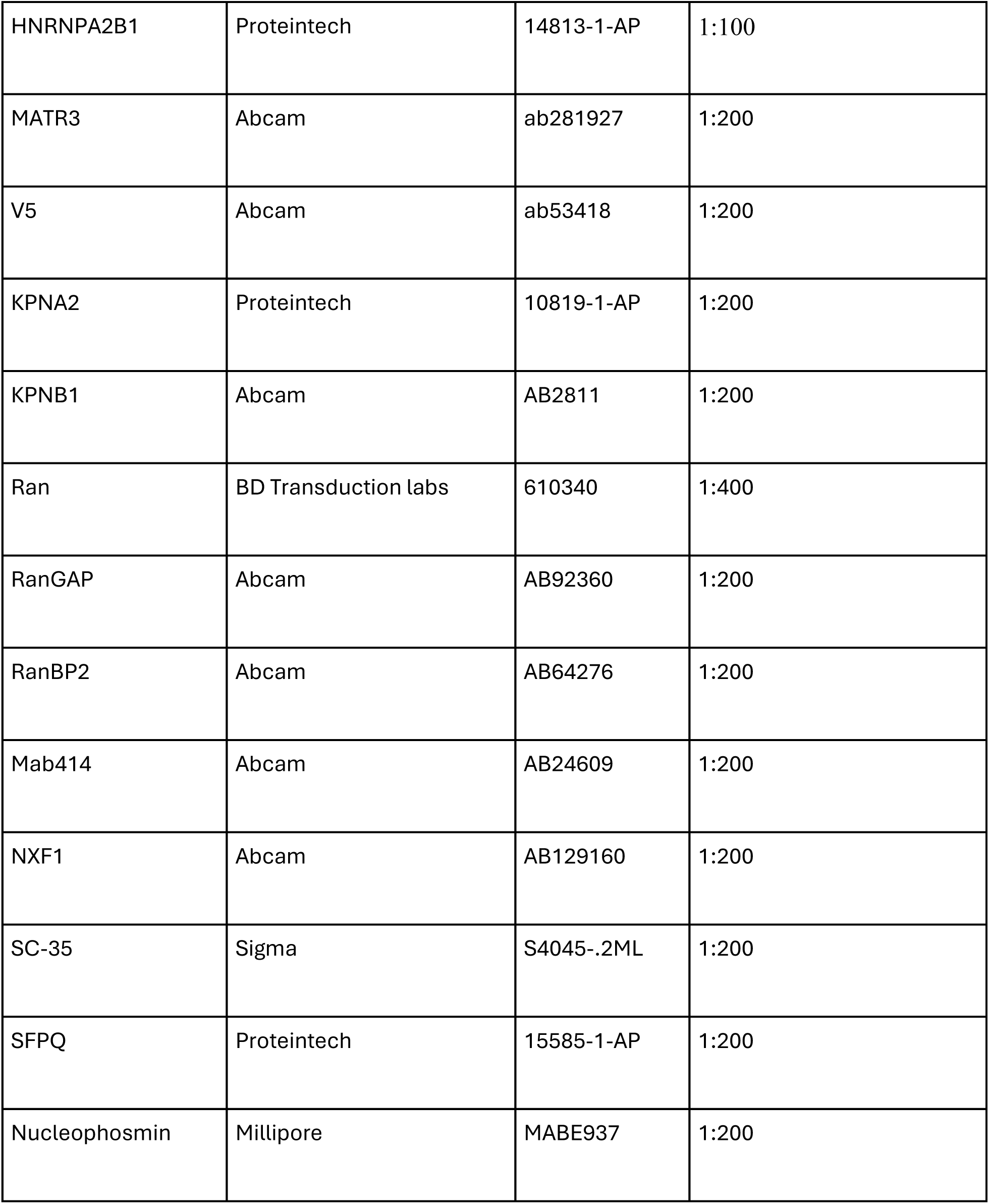

### Plasmids

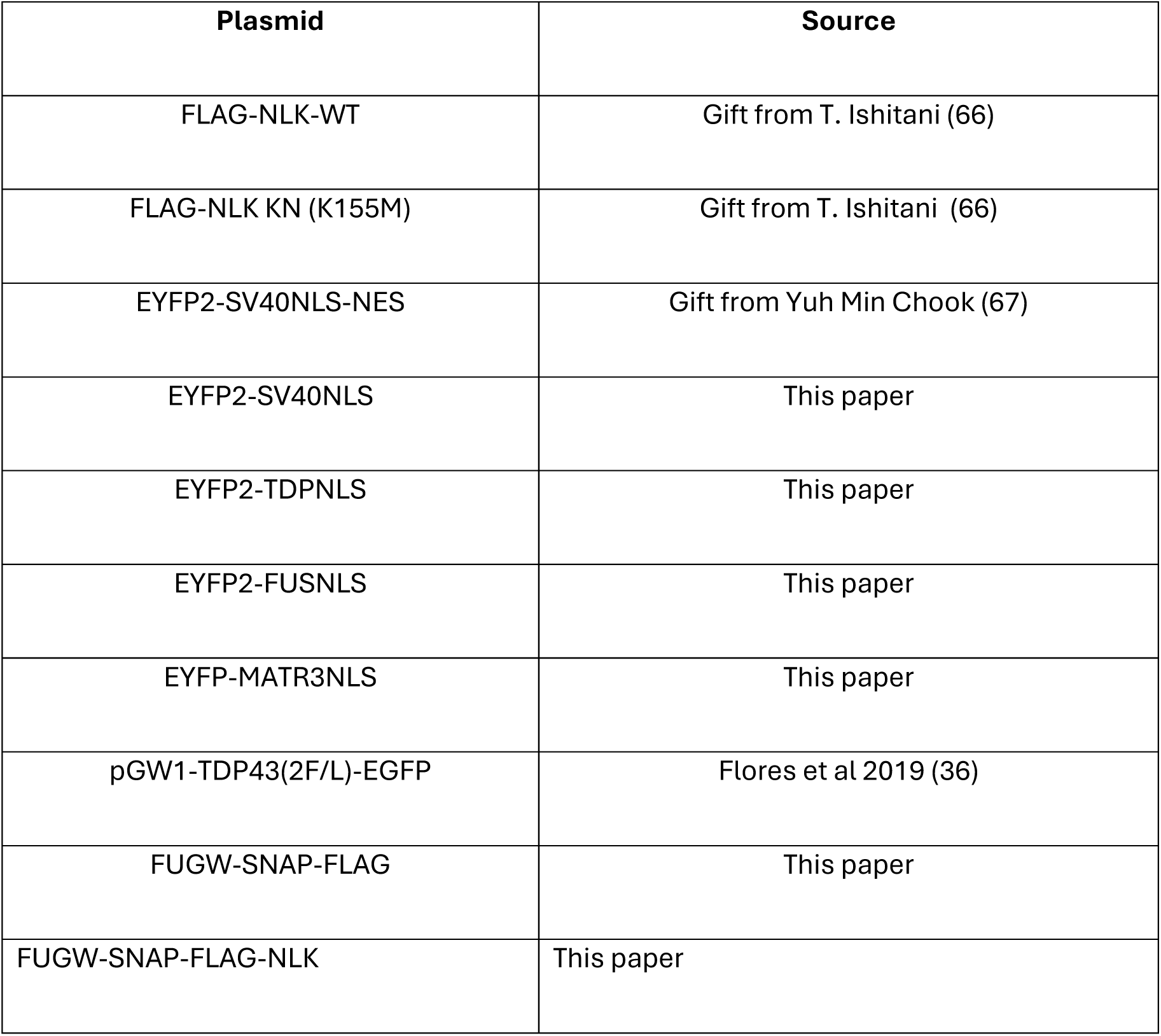

FLAG-NLK-WT and FLAG-NLK KN were a gift from T Ishitani . SNAP-FLAG and SNAP-FLAG NLK were synthesized and subcloned into FUGW. All YFP reporter plasmids were derived from EYFP2-SV40NLS-NES, a kind gift from Yuh Min Chook (67). Site-directed mutagenesis was used to add a stop codon after the SV40NLS. The NLS of TDP43 (residues 82-98) and FUS (residues 495-526) were PCR amplified using primers below, digested with XbaI and BglII, and cloned into the corresponding sites in EYFP2-SV40NLS-NES. The NLS of MATR3 (residues 583-602) was purchased as a Geneblock from Integrated DNA Technologies (IDT), and then digested with XbaI and BglII, and cloned into the corresponding sites in EYFP2-SV40NLS-NES. All plasmids were verified by Sanger sequencing.

### Primers

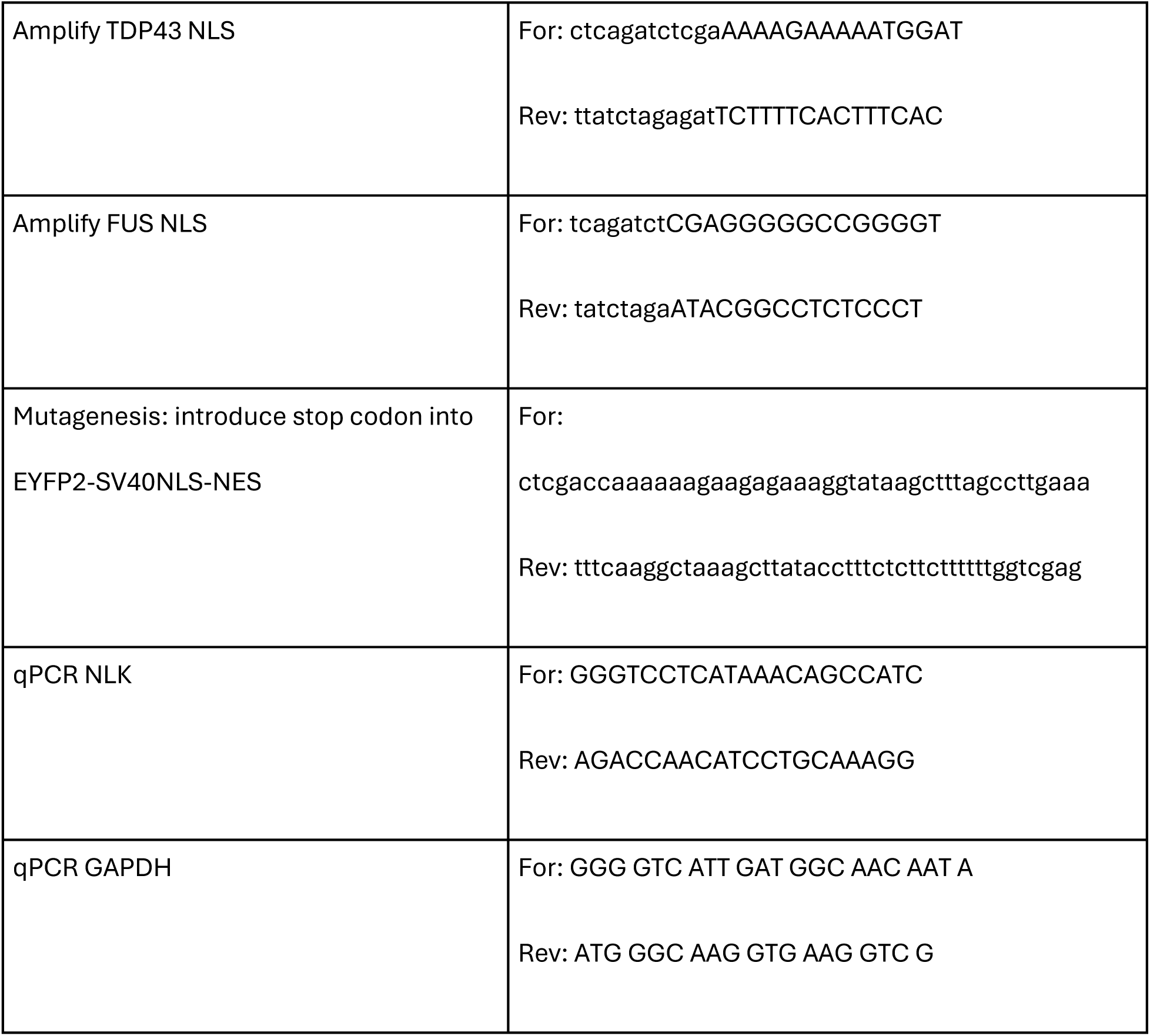

### Data analysis and statistics

Statistical information, including mean and statistical significance values is indicated in the figures or figure legends. At least three biological replicates were used per experiment. For analysis of immunocytochemistry images, ROIs for nucleus and cytoplasm were either generated using CVAT (68) or manually drawn in ImageJ. Nuclear/cytoplasmic ratio was calculated and statistical significance was determined with GraphPadP Prism 9 using the tests indicated in each figure. Data were considered statistically significant at *P < 0.05, **P < 0.01, ***P < 0.001 and ****P < 0.0001. Data from Liu et al. was analyzed with assistance from the TDP43 loss of function database (69); DESeq2 *P*-values were calculated using Wald tests and corrected for multiple testing using the Benjamini–Hochberg method. DEU events with *P*adj <0.05 were considered significantly changed.

### Study approval

Skin samples for iPSC creation were collected and de-identified in collaboration with the Michigan Institute for Clinical and Health Research (MICHR, UL1TR000433) through an institutional review board (IRB)-approved protocol (HUM00028826).

## Data availability

All correspondence regarding the materials and data presented in this paper should be addressed to.

## Author Contributions

MB and SB designed the study. EP and MB designed experiments with SB. MB and EP collected data for most experimental studies, analyzed the data, assembled the figures, and wrote the manuscript. MB and EP performed immunocytochemistry. JM assisted with data collection and analysis. MB performed survival assays in cultured rat neurons and iNeurons, wrote custom FIJI scripts for quantitation of immunocytochemistry and dual IHC. EP made YFP reporter constructs, supervised dual IHC (with the IHC core),and selected motor neurons for quantitation. LGG and MM contributed to mbOrg concept, development, and data analysis. MK and SC cultured mbOrgs and performed RNASeq and qPCR. EU contributed to original concept and development of mbOrgs, and provided RNA-seq data from mbOrgs. XL isolated and cultured rat cortical neurons. EMHT derived iPSC lines and developed protocols for their differentiation. JW developed original code for neuron segmentation and image processing. SB provided resources, funding, and conceptual input for experiments and supervised the research. All authors were involved throughout the research process, agreed amongst themselves regarding roles and responsibilities, and contributed to the review, editing, and approval of the manuscript.

## Acknowledgements

This work was supported by: National Institutes of Health AWD012778 (EP), National Institutes of Health (R01NS097542, R01NS113943 and 1R56NS128110-01 to SJB; R44NS124457 to MM; P30AG072931 to the University of Michigan Brain Bank and Alzheimer’s Disease Research CenterNational Institute on Drug Abuse R44NS124457 (MDM and EMU), the family of Angela Dobson and Lyndon Welch, the A. Alfred Taubman Medical Research Institute, the Danto Family, Ann Arbor Active Against ALS, and the Robert Packard Center for ALS Research.

We thank all members of the Barmada laboratory for their advice and suggestions. We thank Dr. Dafydd Thomas of the Rogel Cancer Center Tissue and Molecular Pathology Shared Resource Laboratory at the University of Michigan (NIH P30 CA04659229). We thank the Michigan Institute for Clinical and Health Research (MICHR, UL1TR000433), who collaborated to collect and de-identify skin samples for iPSC creation. Finally, we thank the patients that donated tissue samples to make this work possible.

Schematics were created in BioRender. Barmada, S. (2024) BioRender.com/z94b294

