## Supplemental Materials for "Nemo-like kinase disrupts nuclear import and drives TDP43 mislocalization in ALS"

#### Supplemental Figures

##### Supplemental Figure 1

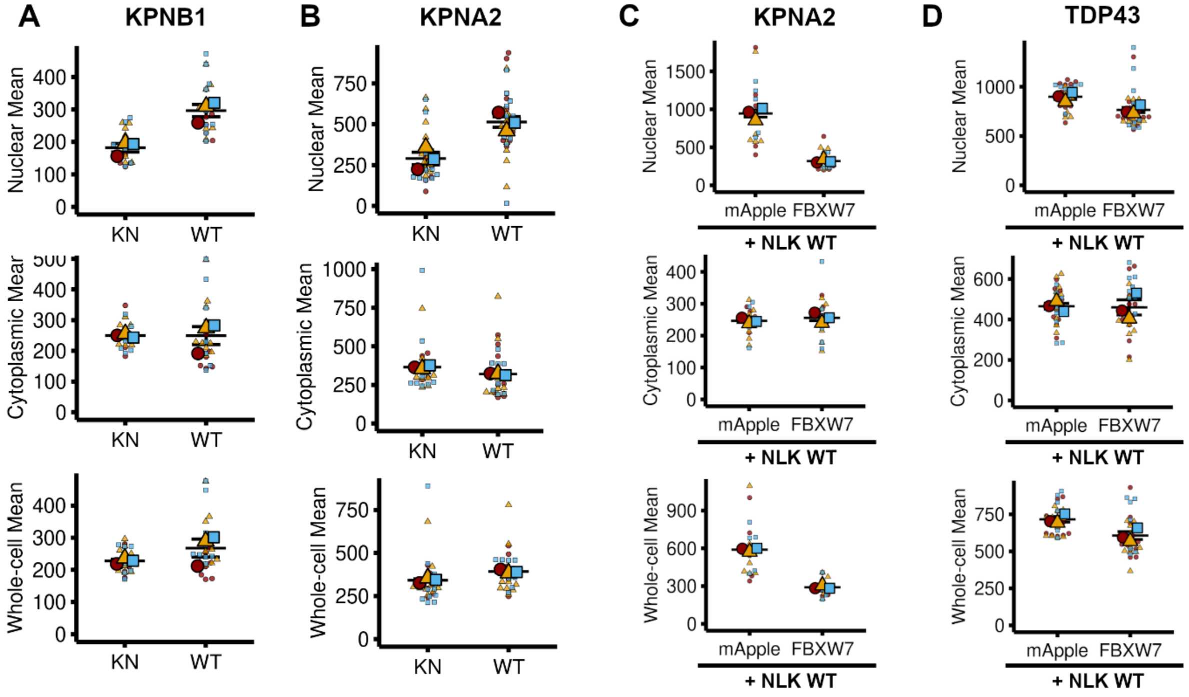


**Supplemental Figure 1. Supporting quantitation from Figure 3.** (**A**) Superplot of KPNB1 Nuclear, Cytoplasm, and Whole-cell levels after NLK KN or NLK WT overexpression related to Figure 3A. Statistical significance indicated as such in all sub-panels: n.s. = not significant, *p<0.05, ** p<0.01, ***p<0.001. (unpaired T-test with Welch's correction). (**B**) Superplot of KPNA2 Nuclear, Cytoplasm, and Whole-cell levels after NLK KN or NLK WT overexpression related to Figure 3B. Line = mean, error bar = standard error. (**C**) Superplot of KPNA1 Nuclear, Cytoplasm, and Whole-cell levels after co-expression of FLAG-NLK WT with either mApple or V5-FBXW7 related to Figure 3C. Line = mean, error bar = standard error. (**D**) Superplot of TDP43 Nuclear, Cytoplasm, and Whole-cell levels after co-expression of FLAG-NLK WT with either mApple or V5-FBXW7 related to Figure 3D. Line = mean, error bar = standard error.

#### Supplemental Figure 2

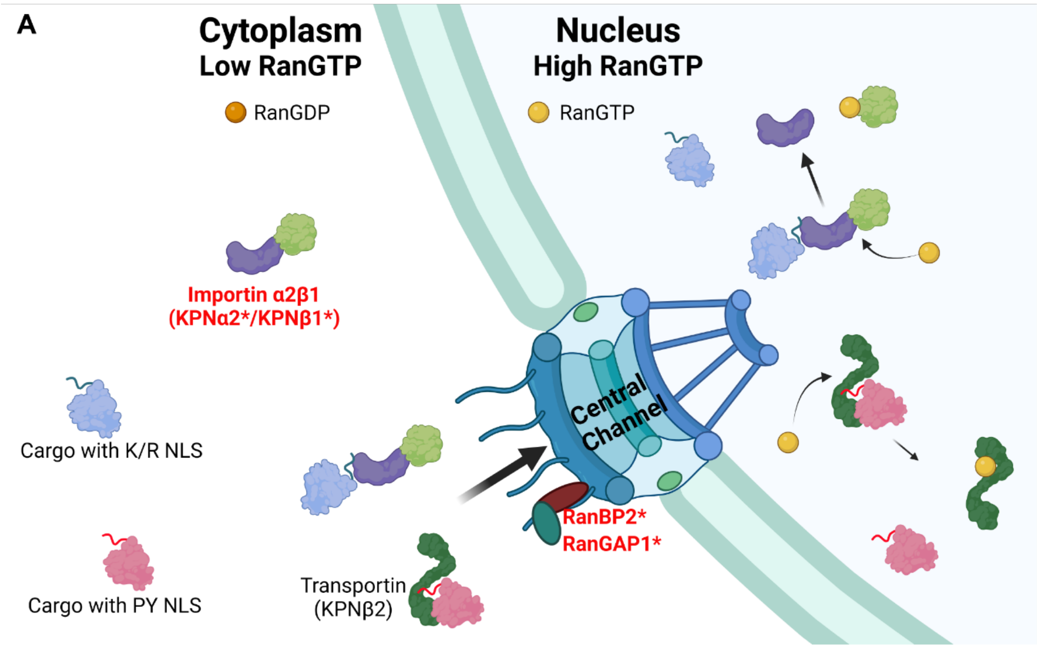


**Supplemental Figure 2. Schematic supporting Figure 4**. (**A**) Schematic of factors involved in nuclear transport. Proteins that directly bind NLK are indicated by red bold typeface and denoted with an asterisk. Schematic created in BioRender.

**Supplemental Figure 3**

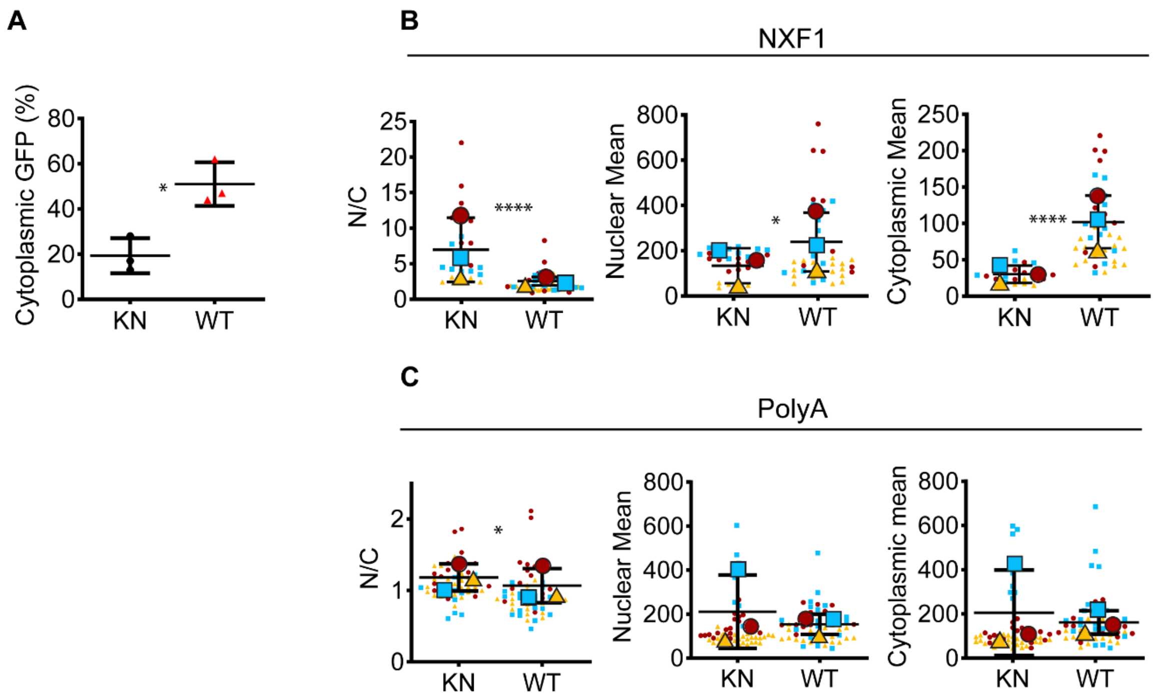


**Supplemental Figure 3. Supporting quantitation from Figure 5.** (**A**) Percentage of cells with cytoplasmic GFP TDP F2L signal in cells expressing FLAG NLK-KN or FLAG NLK-WT, representative images in Fig5A. Bar = mean, error bar = standard deviation. * $p<0.05$ (unpaired t-test with Welch's correction). (**B**) Superplot of nuclear signal, cytoplasmic signal, and N/C ratio of NXF1 in HEK cells expressing FLAG NLK-KN or FLAG NLK-WT, representative images in Fig 5B. Line = mean, error bar = standard deviation. * $p<0.05$, **** $p<0.0001$(unpaired t-test with Welch's correction). (C) Superplot of nuclear signal, cytoplasmic signal, and N/C ratio of PolyA FISH in HEK cells expressing FLAG NLK-KN or FLAG NLK-WT, representative images in Fig 5C. Bar = mean, error bar = standard deviation. * $p<0.05$ (unpaired t-test with Welch's correction).

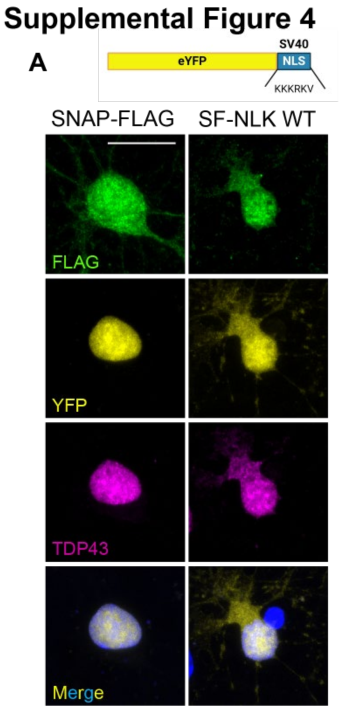


**Supplemental Figure 4. Additional images related to Figure 6.** (**A**) Rodent primary cortical neurons were co-transfected with YFP-NLS$^{SV40}$ and either SNAP-FLAG (SF; negative-control) or SNAP-FLAG-NLK (SF-NLK) followed by immunofluorescence using antibodies against FLAG and TDP43 while DNA was stained with Hoechst. Scale bar = 10 μm.

### Supplemental Figure 5

**A**

| | ALS - 1 | ALS - 2 | ALS - 3 | ALS - 4 | Ctrl - 1 | Ctrl - 2 | Ctrl - 3 | Ctrl - 4 |
|---|---|---|---|---|---|---|---|---|
| Male/female (M/F) | M | F | M | M | M | F | M | F |
| Age (y) | 64 | 76 | 64 | 51 | 61 | 78 | 72 | 64 |
| Clinical diagnosis | ALS with FTLD | ALS | ALS | ALS | Early-onset AD | Probable AD | Lewy Body Dementia | HD |
| Neuropathological diagnosis | ALS<br>PSP | ALS | ALS | ALS | ADNC, high | ADNC, int | ADNC, high<br>LATE-NC, stage 1 | HD<br>ADNC, low |

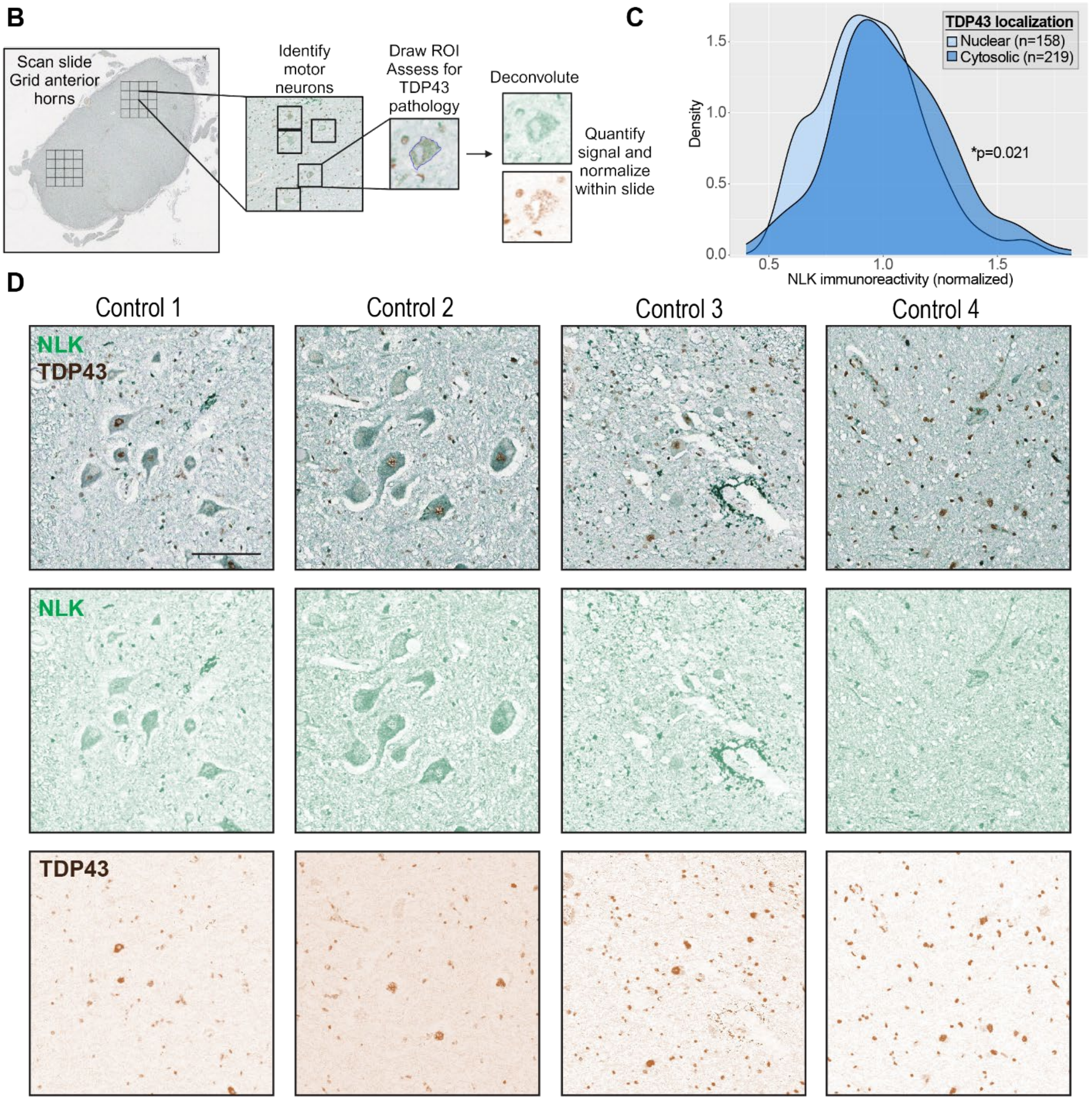

**Supplemental Figure 5. NLK/TDP43 dual immunohistochemistry in patient tissue.** (**A**) Patient characteristics. Ctrl- Control. ALS- Amyotrophic lateral sclerosis. HD- Huntington's Disease. AD- Alzheimer's Disease. ADNC- Alzheimer Disease Neuropathologic change. LATE-NC- Limbic predominant age-related TDP43 encephalopathy neuropathologic change. (**B**) Schematic showing workflow for quantification of dual immunohistochemistry. (**C**) Density plot depicting the change in NLK immunoreactivity in motor neurons with and without TDP43 pathology. * $p<0.05$ by 2-sided Kolmogorov Smirnov test. Representative images in 7E. (**D**) Dual immunohistochemistry for NLK and TDP43, performed on spinal cord tissue from four control patients without spinal cord pathology. Scale bar = 100 μm.

#### Supplemental Figure 6

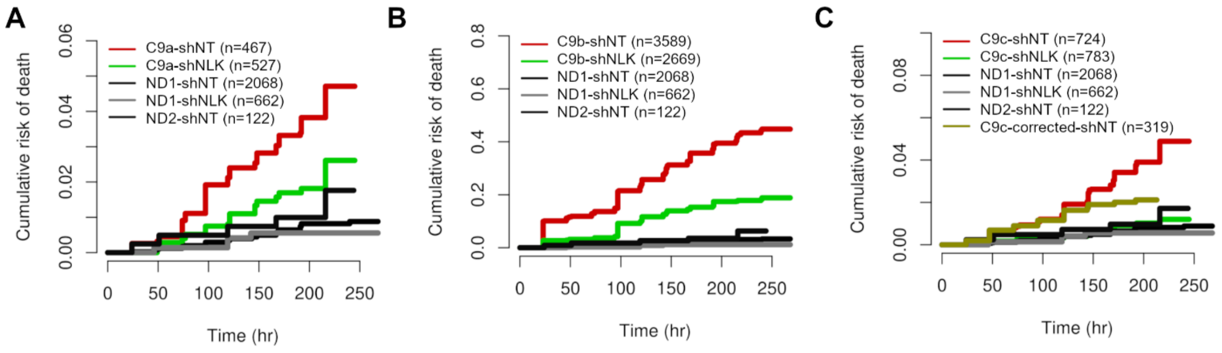


**Supplemental Figure 6. Supporting quantification for Figure 7.** (**A**) Cumulative hazard plot showing relative risk of death in neurons from a single patient-derived C9orf72 iPSC line (C9a) after transduction with lentivirus encoding either Scramble of NLK shRNA compared to two independent non-disease (ND) lines. (**B**) Cumulative hazard plot showing relative risk of death in neurons from a single patient-derived C9orf72 iPSC line (C9b) after transduction with lentivirus encoding either Scramble of NLK shRNA compared to two independent non-disease (ND) lines. (**C**) Cumulative Hazard plot showing relative risk of death in neurons from a single patient-derived C9orf72 iPSC line (C9c) after transduction with lentivirus encoding either Scramble of NLK shRNA compared to two independent non-disease (ND) lines.

**Supplemental Methods:**

**iPSC lines:**

| | **ID** | **Onset age** | **Biopsy age** | **Gender** | **Onset** | **Notes** |
|---|---|---|---|---|---|---|
| **C9orf72-fALS** | C9b (883) | 49 | 51 | M | Lumbar | C9orf72 mutation + |
| | C9a (312) | 52 | 54 | M | Lumbar | C9orf72 mutation + |
| | C9c (CS52) | 46 | 47 | M | Lumbar | C9orf72 mutation + |
| **TDP43-fALS** | TDP43 (M337V) | | 54 | F | | M337V and C-terminal Dendra2 inserted into ND1 (1021) by CRISPR/Cas9 |
| | TDP43 (WT) | | 54 | F | | C-terminal Dendra2 inserted intoND1 (1021) by CRISPR/Cas9 |
| **Control** | ND1 (1021) | | 54 | F | | Healthy |
| | ND2 (746) | | 58 | M | | Healthy |
| | C9c-corrected | 46 | 47 | M | Lumbar | C9orf72 mutation removed from CS52i by CRISPR/Cas9 |

**Antibodies:**

| | Vendor | Catalog # | Concentration for ICC |
|---|---|---|---|
| Flag | AbCam | ab1170 | 1:200 |
| TDP43 | R&D | MAB7778 | 1:200 |
| TDP43 | Proteintech | 10782-2-AP | 1:200 |
| FUS | Sigma | HPA008784 | 1:200 |
| HNRNPA2B1 | Proteintech | 14813-1-AP | 1:100 |
| MATR3 | Abcam | ab281927 | 1:200 |
| V5 | Abcam | ab53418 | 1:200 |
| KPNA2 | Proteintech | 10819-1-AP | 1:200 |
| KPNB1 | Abcam | AB2811 | 1:200 |
| Ran | BD Transduction labs | 610340 | 1:400 |
| RanGAP | Abcam | AB92360 | 1:200 |
| RanBP2 | Abcam | AB64276 | 1:200 |
| Mab414 | Abcam | AB24609 | 1:200 |

| | | | |
|---|---|---|---|
| NXF1 | Abcam | AB129160 | 1:200 |
| SC-35 | Sigma | S4045-.2ML | 1:200 |
| SFPQ | Proteintech | 15585-1-AP | 1:200 |
| Nucleophosmin | Millipore | MABE937 | 1:200 |

**Plasmids**

| Plasmid | Source |
|---|---|
| FLAG-NLK-WT | Gift from T. Ishitani (66) |
| FLAG-NLK KN (K155M) | Gift from T. Ishitani (66) |
| EYFP2-SV40NLS-NES | Gift from Yuh Min Chook (67) |
| EYFP2-SV40NLS | This paper |
| EYFP2-TDPNLS | This paper |
| EYFP2-FUSNLS | This paper |
| EYFP-MATR3NLS | This paper |
| pGW1-TDP43(2F/L)-EGFP | Flores et al 2019 (36) |
| FUGW-SNAP-FLAG | This paper |
| FUGW-SNAP-FLAG-NLK | This paper |

**Primers**

| | |
|---|---|
| Amplify TDP43 NLS | For: ctcagatctcgaAAAAGAAAAATGGAT<br><br>Rev: ttatctagagatTCTTTTCACTTTCAC |
| Amplify FUS NLS | For: tcagatctCGAGGGGGCCGGGGT<br><br>Rev: tatctagaATACGGCCTCTCCCT |
| Mutagenesis: introduce stop codon into EYFP2-SV40NLS-NES | For: ctcgaccaaaaaagaagagaaaggtataagctttagccttgaaa<br><br>Rev: tttcaaggctaaagcttatacctttctcttcttttttggtcgag |
| qPCR NLK | For: GGGTCCTCATAAACAGCCATC<br><br>Rev: AGACCAACATCCTGCAAAGG |
| qPCR GAPDH | For: GGG GTC ATT GAT GGC AAC AAT A<br><br>Rev: ATG GGC AAG GTG AAG GTC G |
